# Dynamic *in situ* confinement triggers ligand-free neuropeptide receptor signaling

**DOI:** 10.1101/2021.12.15.472742

**Authors:** M. Florencia Sánchez, Marina S. Dietz, Ulrike Müller, Julian Weghuber, Karl Gatterdam, Ralph Wieneke, Mike Heilemann, Peter Lanzerstorfer, Robert Tampé

## Abstract

Membrane receptors are central to cell-cell communication. Receptor clustering at the plasma membrane modulates physiological responses, and mesoscale receptor organization is critical for downstream signaling. Spatially restricted cluster formation of the neuropeptide Y_2_ hormone receptor (Y_2_R) was observed *in vivo*; however, the relevance of this confinement is not fully understood. Here, we controlled Y_2_R clustering *in situ* by a chelator nanotool. Due to the multivalent interaction, we observed a dynamic exchange in the microscale confined regions. Fast Y_2_R enrichment in clustered areas triggered a ligand-independent downstream signaling determined by an increase in cytosolic calcium, cell spreading, and migration. We revealed that the cell response to ligand-induced activation was amplified when cells were pre-clustered by the nanotool. Ligand-independent signaling by clustering differed from ligand-induced activation in the binding of arrestin-3 as downstream effector, which was recruited to the confined regions only in the presence of the ligand. This approach enables *in situ* clustering of membrane receptors and raises the possibility to explore different modalities of receptor activation.

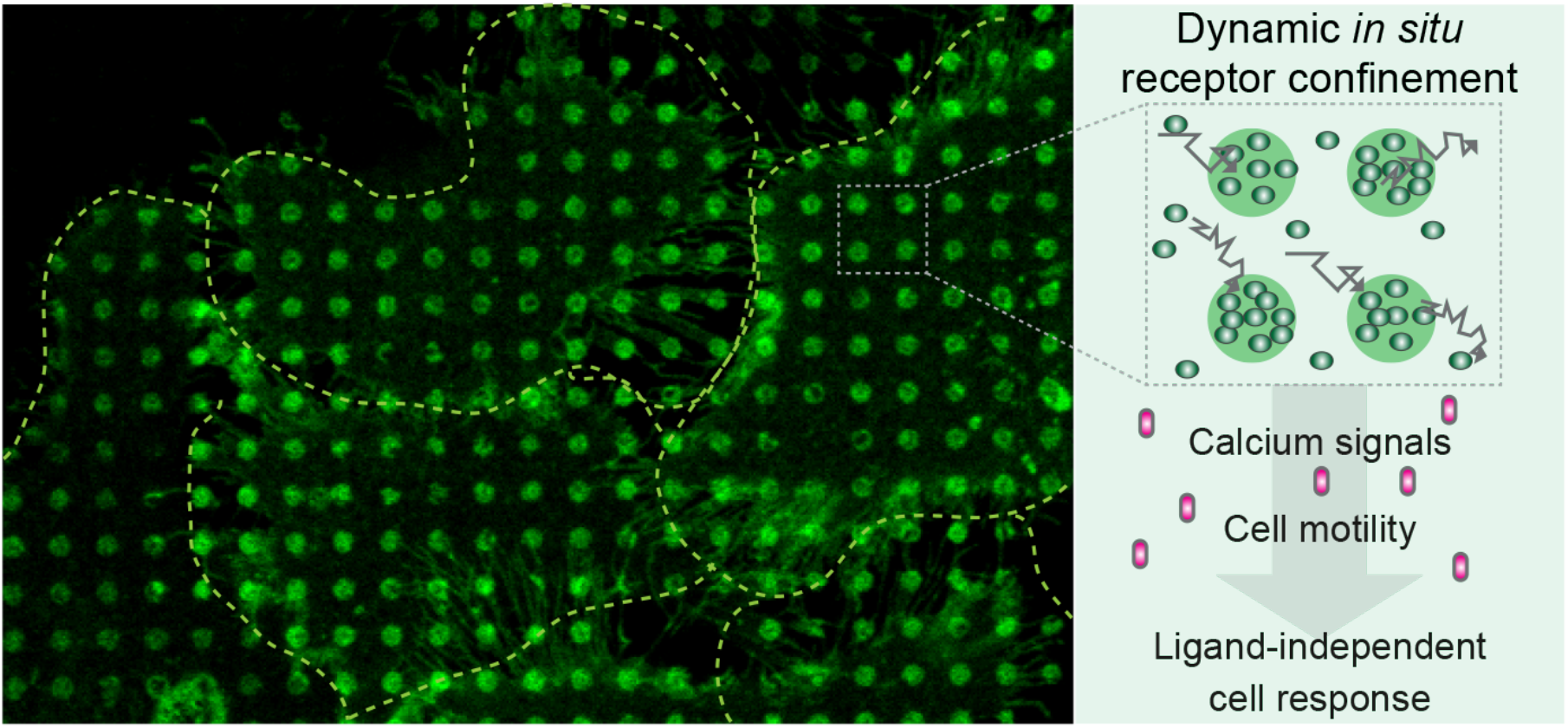

## Introduction

Cells translate stimuli into biochemical signals through membrane receptors controlling multiple aspects of cell behavior, including migration (Kupperman *et al*., 2000; Stallaert *et al*., 2018), differentiation (Li & Rudensky, 2016; Luther & Cyster, 2001), apoptosis (Scott *et al*., 2009), as well as infectious diseases and cancer (Boncompain *et al*., 2019; Haqshenas & Doerig, 2019; Kawai & Akira, 2005; Pasquale, 2010; Pike *et al*., 2018; Sebestyen *et al*., 2020; Tsukiyama *et al*., 2020). Receptors form dynamic assemblies or clusters that modulate downstream signaling and the final physiological response. Upon activation, these receptors undergo transitions from freely diffusing monomers to less mobile nanoclusters and further to higher-order oligomers (Ojosnegros *et al*., 2017; Su *et al*., 2016). In signal transduction, the mechanisms for receptor cluster formation and cluster behavior have become physiologically relevant topics. However, the role of mesoscale (hundreds of nanometers) receptor organization in signal transduction remains unsolved, mainly because techniques to trigger receptor clustering *in situ* and monitor this assembly process in real-time are largely limited.

Nano- and microlithographic approaches have provided cell-compatible scaffolds to investigate confined ligand-receptor interactions. Various techniques, ranging from photolithography (Chen *et al*., 2021; Scheideler *et al*., 2020; Traub *et al*., 2016) to electron-beam lithography (Cai *et al*., 2018; Nassereddine *et al*., 2021) and microcontact printing (μCP) (Lindner *et al*., 2018; Sánchez *et al*., 2018), have yielded information on how topology and mobility of the stimulus regulate cellular outcomes. Recently, optogenetics and optochemistry have provided the possibility of targeting receptor oligomerization with high spatiotemporal control (Bardhan & Deiters, 2019; Goglia & Toettcher, 2019; Taslimi *et al*., 2014). However, approaches that can be easily adapted to a variety of receptors or experimental setups and that offer the ability to analyze large cell populations simultaneously are rare.

Heterotrimeric guanine nucleotide-binding protein (G protein)-coupled receptors (GPCR) are key cell surface proteins that regulate a plethora of cellular responses to external stimuli (Hilger *et al*., 2018; Venkatakrishnan *et al*., 2013; Wootten *et al*., 2018). The Y_2_ receptor (Y_2_R) is one of the four human neuropeptide Y (NPY) receptor subtypes, which belong to the rhodopsin-like (class A) GPCR superfamily (Parker & Balasubramaniam, 2008; Tang *et al*., 2022). Y_2_R is linked to many important physiological processes, such as fear extinction (Méndez-Couz *et al*., 2021), regulation of food intake (Huang *et al*., 2014), and obesity (Lafferty *et al*., 2021). Y_2_R activation by neuropeptide Y (NPY) has been shown to promote cell migration and proliferation (Ekstrand *et al*., 2003; Movafagh *et al*., 2006). It has been recently demonstrated that Y_2_Rs respond to light-guided microscale clustering at spatially defined locations (Sánchez *et al*., 2021). Y_2_Rs are activated independently of canonical ligands, evoking elevated cytosolic calcium, a change in cell spreading behavior, and a localized migratory pattern.

Here, we established a versatile approach for *in situ* receptor clustering using a multivalent chelator nanotool (*tris N*-nitrilotriacetic acid, *tris*NTA), which displays high affinity for histidine (His)-tagged proteins. The nanometer size of the tool in combination with pre-structured matrices enabled receptor clustering with high spatiotemporal resolution. The lateral organization of Y_2_Rs in living cells was controlled within minutes in a non-invasive and ligand-independent manner. Microscale receptor clusters with a high degree of homogeneity in size and density were generated at the plasma membrane. Analysis of the receptor mobility revealed a dynamic assembly with fast exchange of the receptors within the confined areas in contrast to the static clustering induced by an anti-His-tag antibody on the same matrix. Nanotool-induced receptor clustering triggered ligand-independent activation of signal transduction, as evidenced by an increase in cytosolic calcium and cell motility, effects also observed in ligand-induced receptor activation. Moreover, we demonstrated an amplification of the signal upon ligand-induced activation in cells pre-clustered with the nanotool. As additional downstream event, we uncovered high arrestin-3 (Arr3) co-recruitment to the patterned areas only in the presence of the canonical ligand, suggesting an Arr3-independent desensitization mechanism for the ligand-independent response. Compared to standard micropatterning techniques, this generic approach advances ligand-free receptor signaling studies, with the advantage that large cell populations can be imaged simultaneously, and no expensive equipment is required for implementation. The versatile nanotool can be adapted to a variety of systems and receptors through minimal modifications.

## Results

### *In situ* receptor clustering by a multivalent nanotool

We developed a system to induce dynamic receptor assembly *in situ* based on a multivalent chelator *tris*NTA nanotool (**Figure 1A**), which is equipped with a biotin moiety (*tris*NTA^PEG12-B^) (**Figure 1B**). *tris*NTA^PEG12-B^ displays a high affinity for His_6_-tagged proteins (*K*_D_ ≈ 1-10 nM), resulting in a site-specific but reversible interaction with minimal steric constraints (Gatterdam *et al*., 2018). Microcontact printing is a widely used method to investigate protein-protein interactions in living cells (Ruiz & Chen, 2007; Torres *et al*., 2008). However, reproducible patterned substrates with a generic structure over extensive millimeter dimensions which allow simultaneous analyses of large cell populations are difficult to produce. We used a large-area perfluoropolyether (PFPE) elastomeric stamps inked with bovine serum albumin (BSA) to print 96-well size glass (Hager *et al*., 2021; Lanzerstorfer *et al*., 2020). Wells within these plates 104 containing a BSA-structured matrix were functionalized with biotinylated BSA (biotin-BSA) and 105 streptavidin (SA) (**Figure 1A**). Subsequent functionalization with the nanotool and His_6_-tagged fluorescent proteins resulted in well-resolved protein patterns that were analyzed by confocal laser scanning microscopy (CLSM). The results confirmed the specificity of the nickel-loaded *tris*NTA chelator to capture His_6_-tagged proteins in defined regions of 1 μm or 3 μm diameter (**Figure 1C, D**).

**Figure 1.**
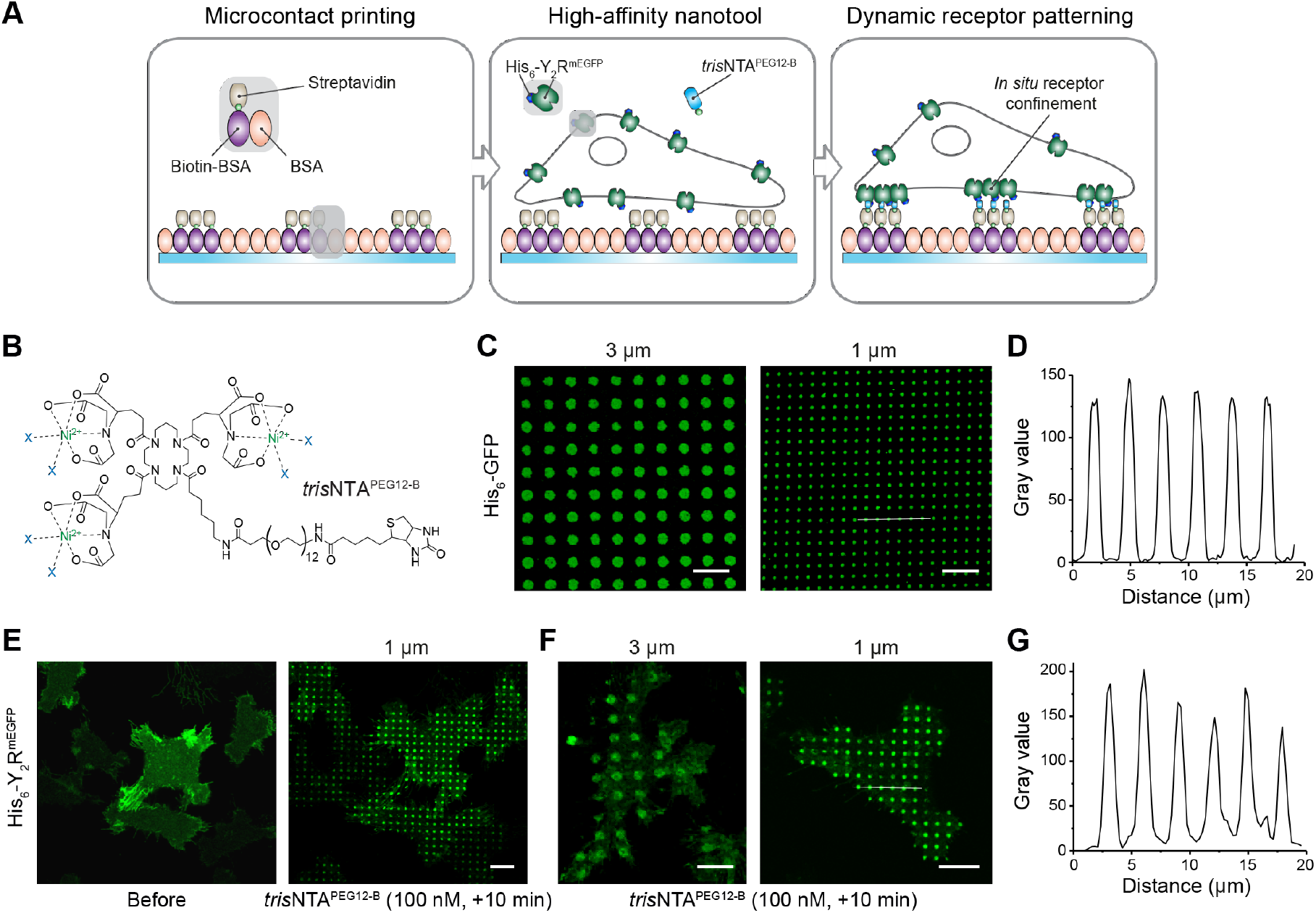
*In situ* ligand-free receptor confinement. (A) Rational of the experimental design for ligand-free receptor clustering. Matrices pre-structured with BSA are stepwise functionalized with biotin-BSA and SA. Upon addition of the multivalent nanotool *tris*NTA^PEG12-B^, His_6_-tagged receptors in HeLa cells are captured to the pre-structured regions via multivalent His-tag/*tris*NTA interaction. (B) Chemical structure of the *tris*NTA^PEG12-B^. (C) Variable size protein patterns generated by further functionalization of SA-matrices with the nanotool followed by incubation with His_6_-GFP (0.1 μM, 20 min). Images were acquired by confocal laser scanning microscopy (CLSM). (D) Intensity profile of the 1 μm pattern (white line in (C)) reflects high specificity of the interaction. (E) Large-scale cell patterning in living cells occurred 10 min after incubation with the nanotool (*tris*NTA^PEG12-B^ 100 nM final, 10 min). Y_2_R-expressing cells were allowed to adhere to the functionalized matrix for 3 h and immediately imaged by CLSM in live-cell imaging solution (LCIS) at 37 °C. (F) Customized Y_2_R assembly on 3 μm and 1 μm SA-pre-structured matrices. (G) Intensity profile of the 1 μm pattern (white line in (F)) showed an intensity comparable to a soluble His_6_-tagged protein. Scale bars: 10 μm.

The ability to control the organization of membrane receptors *in situ* is important for dissecting the spatial complexity of cell signaling and the extracellular environment. With this aim, we established a monoclonal human cervical cancer HeLa cell line expressing low amounts of Y_2_R (∼300,000 receptors/cell) utilizing a tetracycline-inducible (T-Rex) expression system (Sánchez *et al*., 2021). Y_2_R displayed an N-terminal His_6_-tag to the extracellular space and a cytosolic C-terminal monomeric Enhanced Green Fluorescent Protein (mEGFP) (His_6_-Y_2_R^mEGFP^, in brief Y_2_R). It is important to mention that these modifications do not affect receptor activity, selectivity, or ligand binding as previously shown (Sánchez *et al*., 2021). It has been further demonstrated that Y_2_R does not require the N terminus for ligand binding (Lindner *et al*., 2009). Y_2_R-positive cells properly adhered to 1 μm and 3 μm SA-functionalized matrices and showed a homogeneous receptor distribution at the basal plasma membrane (**Figure 1E**). Addition of *tris*NTA^PEG12-B^ (100 nM final) triggered receptor assembly. Within five minutes, all cells showed receptor patterns at the plasma membrane comparable in size and density (**Figure 1E, F, Figure 1–figure supplement 1**). Importantly, recruitment of soluble His_6_-tagged GFP proteins as well as Y_2_Rs to 1 μm pre-structured spots led to analogous intensity profiles, reflecting that similar densities were obtained in both cases (**Figure 1D, G**).

In contrast, cells expressing Y_2_Rs without the His_6_-tag (Y_2_R^mEGFP^) showed no receptor clustering after addition of *tris*NTA^PEG12-B^ (**Figure 1–figure supplement 2**), demonstrating the specificity of the His_6_-tag/*tris*NTA interaction. Remarkably, ten minutes after receptor clustering by the multivalent nanotool, the Y_2_R enrichment resulted in an integrated receptor density equivalent to that of cells cultured on matrices functionalized with anti-His_6_ antibodies. However, a 10-fold higher antibody concentration (1 μM final) was required compared to the multivalent nanotool, demonstrating its efficacy in capturing His_6_-tagged Y_2_ receptors (**Figure 1–figure supplement 3**). The nanotool-induced 3 μm clusters presented a 9-fold increase in integrated density compared to 1 μm arrays, consistent with the increase in pattern area. Overall, our approach enabled versatile *in situ* receptor clustering with high specificity.

### Receptor diffusion and dynamic exchange in the confined regions

GPCR signaling results from dynamic interactions among receptors, G proteins, and the complex surrounding membrane environment, which confers flexibility and versatility on this fundamental biological process. To characterize receptor clustering induced by the chelator nanotool, we first examined whether Y_2_R clustering affects lipid diffusion and distribution by labeling the membrane with the lipid-like dye CellMask. We observed a homogeneous staining of the plasma membrane, demonstrating that receptor confinement does not affect lipid distribution (**Figure 2–figure supplement 1**). To determine lateral diffusion coefficients (*D*), we performed fluorescence recovery after photobleaching (FRAP). *In situ* receptor clustering was triggered on Y_2_R-expressing cells cultured on SA-matrices by incubation with *tris*NTA^PEG12-B^ (100 nM, +10 min), followed by membrane labeling with the lipid-like dye. In a subsequent step, square-shaped regions of interest (ROIs) covering four 1 μm-sized spots were photobleached. Fluorescence recovery was analyzed by a FRAP simulation approach that enabled calculation of diffusion coefficients independent of bleaching geometry (Blumenthal *et al*., 2015). The lateral diffusion coefficient of lipids obtained by FRAP showed an average value of *D*_lipid_ = 0.66 ± 0.10 μm^2^/s, which is in agreement with literature values for free Brownian lipid diffusion at the plasma membrane (Schwille *et al*., 1999; Wawrezinieck *et al*., 2005).

**Figure 2.**
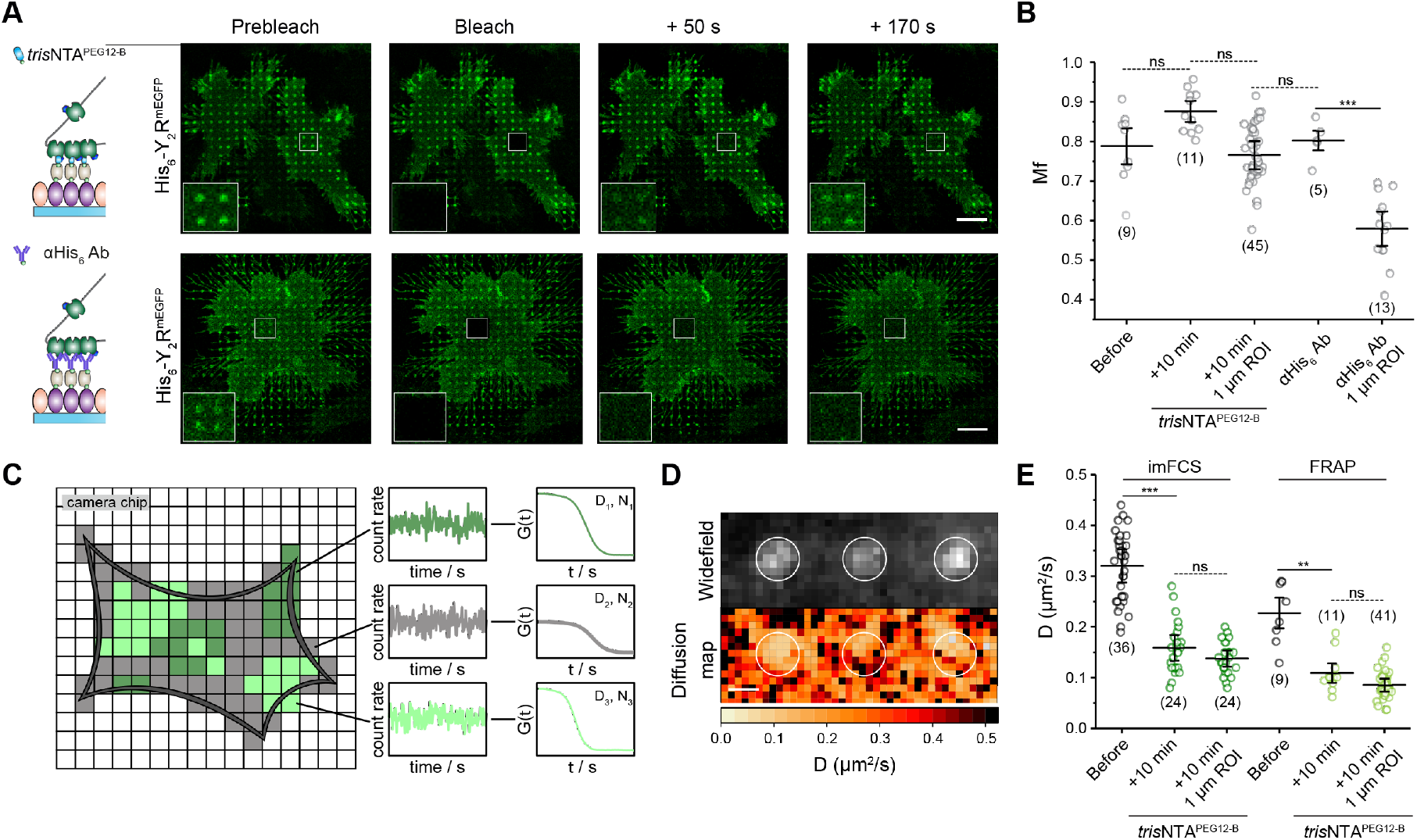
Decrease of receptor mobility in confined regions. (A) FRAP analyses upon Y_2_R clustering induced either by the nanotool *in situ* or by an anti-His_6_ antibody (αHis_6_ Ab). Y_2_R-expressing cells were allowed to adhere to SA- or - αHis_6_ Ab matrices for 3 h and immediately imaged by CLSM in live-cell imaging solution (LCIS) at 37 °C. The *tris*NTA^PEG12-B^ nanotool was added to a final concentration of 100 nM. Insets represent the bleached ROIs. Fast recovery of the clusters can be detected for the case of the multivalent nanotool. (B) Quantification of the receptor mobile fraction for cell patterning by the *tris*NTA^PEG12-B^ and anti-His_6_ antibody demonstrated unchanged receptor mobile fraction for the nanotool, suggesting a high receptor exchange. The mean ± SD is shown. 9 cells before, 11 cells after *tris*NTA^PEG12-B^ addition (45x 1 μm ROIs), and 5 cells on anti-His_6_ antibody matrices (13x 1 μm ROIs) were analyzed. ***p≤ 0.001 for Tukey test. (C) imFCS correlates fluorescence intensity fluctuations in single camera pixels with a large degree of statistics, providing accurate diffusion coefficients with high spatial and temporal resolution. (D) Widefield image of a ROI at the plasma membrane of a living cell upon addition of the nanotool analyzed by imFCS (left). The analyses of numerous pixels simultaneously provide two-dimensional diffusion data that draw a picture of the mobility of membrane receptors and reveal local differences in the diffusion (right). (E) Both techniques demonstrated a decrease in the lateral diffusion of the receptor at the plasma membrane after addition of the chelator nanotool. Analysis of 1 μm clusters within the entire ROI led to a further decrease in the lateral diffusion coefficient. For imFCS analyses, two-sample t-tests (α = 0.05) were applied to compare the diffusion coefficients for the different conditions. The mean ± SD is shown. 36 and 24 cells for the conditions before and after addition of *tris*NTA^PEG12-B^ were analyzed. For FRAP, the mean ± SD is shown. 9 cells before, 11 cells after *tris*NTA^PEG12-B^ addition (41x 1 μm ROIs) were analyzed. ***p≤ 0.001 for Tukey test. Scale bar: 10 μm (A), 1 μm (D).

To evaluate the Y_2_R mobility in the ligand-free induced clusters, we determined the lateral diffusion of Y_2_ receptor in cells cultured on SA-matrices before and after receptor clustering by *tris*NTA^PEG12-B^ (100 nM, 10 min). A square-shaped ROI covering four 1 μm-sized spots was photobleached. A significant decrease in the lateral diffusion of the Y_2_R was observed at the basal membrane of cells after receptor confinement by *tris*NTA^PEG12-B^ (*D*_before_ = 0.25 ± 0.08 μm^2^/s *versus D*_after_ = 0.10 ± 0.03 μm^2^/s) (**Figure 2A, E**). Surprisingly, the receptor intensity showed a high recovery within ∼3 min after photobleaching (**Figure 2A, Figure 2– figure supplement 2, Video 1**). Notably, no significant difference in receptor mobile fraction (*M*_f_) before and after addition of the nanotool was observed (*M*_*f*_ = 0.80 ± 0.04) (**Figure 2B**). In comparison, FRAP analyses of cells cultured on matrices functionalized with anti-His_6_ antibodies presented a drastic decrease in receptor diffusion and mobile fraction at the clustered spots (*M*_f,anti-His6 Ab_ = 0.56 ± 0.08) (**Figure 2A, B, Figure 2–figure supplement 3, Video 2**). Despite the high affinity and kinetically stable binding (*k*_off_ = 0.18 h^-1^) (Gatterdam *et al*., 2018), the His-tag/*tris*NTA system relies on molecular multivalency, which enables competition of binding sites with histidine or other receptors, thus making the process of receptor assembly reversible. We rationalized that free receptors diffuse into the clustered spots and exchange with photobleached receptors at multivalent binding sites, leading to a dynamic confinement. Our results indicate that a high proportion of receptors is exchanged in and out of micrometer-sized clusters, an effect that likely depends on cluster size, with larger clusters showing less recovery (Sánchez *et al*., 2021).

We also investigated the lateral receptor mobility with a higher spatiotemporal resolution using imaging fluorescence correlation spectroscopy (imFCS). FCS is used to study the diffusion of membrane proteins in living cells with single-molecule sensitivity (**Figure 2C**). These multiplexed FCS measurements are realized by analyzing many pixels simultaneously using a widefield setup (Harwardt *et al*., 2018; Kannan *et al*., 2006). Regions of interests (ROIs) on Y_2_R-expressing cells cultured on SA-matrices were analyzed before and after receptor clustering by *tris*NTA^PEG12-B^. Enrichment of Y_2_R at the basal membrane was observed with total internal reflection fluorescence (TIRF) microscopy (**Figure 2D**). Consistent with the FRAP measurements, the Y_2_R diffusion coefficient decreased upon cluster formation (*D*_before_ = 0.32 ± 0.06 μm^2^/s and *D*_after_ = 0.16 ± 0.05 μm^2^/s). The receptor diffusion coefficient measured before clustering was comparable to membrane proteins of similar size (Lippincott-Schwartz *et al*., 2001), demonstrating that the microstructured confinement does not affect receptor mobility.

In contrast to FRAP, imFCS provides a two-dimensional diffusion map, which enables the determination of local differences in the lateral diffusion coefficient of membrane receptors with high precision. Quantitative analysis of the 1 μm cluster spots in the acquired ROIs resulted in a lateral diffusion coefficient of *D*_spots_ = 0.14 ± 0.03 μm^2^/s (**Figure 2E**). Taking into consideration that imFCS detects mobile particles only, we determined a similar decrease in lateral diffusion in the patterned regions for cells cultured on matrices functionalized with anti-His_6_ antibodies (**Figure 2–figure supplement 3**). Taken together, we unravel that in microscale clusters, associations between His_6_-tagged Y_2_Rs and multivalent *tris*NTA^PEG12-B^ resulted in a decreased lateral diffusion but dynamic receptor exchange with unchanged mobile fraction, which is similar to the behavior described for ligand-activated receptor clustering (Chavez-Abiega *et al*., 2019).

### Ligand-independent receptor clustering triggers fast signaling

To mimic a scenario in which receptors can cluster at the mesoscale and still reflect physiologically relevant dimensions for clustering at cell-cell interfaces (Guo *et al*., 2008), Y_2_R-expressing cells were cultured on microstructured matrices with a diameter of 1 μm, the smallest pattern we can produce and analyze with high accuracy. After addition of the multivalent nanotool, the receptor redistribution was tracked by CLSM at 37 °C. Receptor clustering occurred in the first minutes and increased within 10 min until an equilibrium was reached, resulting in a 2.5-fold increase in receptor density compared to the initial state (**Figure 3A, Video 3**). The kinetic profile of Y_2_R recruitment to the 1 μm spots followed a pseudo-first-order assembly rate of 0.35 ± 0.05 min^-1^ (**Figure 3A, B**). Considering the average cell area of 1,420 ± 50 μm^2^ (n = 66 cells) and the enrichment factor (2.5-fold), we estimated a receptor density of ∼500 receptors/μm^2^ in the patterned regions (∼400 receptors per 1 μm circular spot), a value comparable to other receptor studies utilizing fluorescence correlation spectroscopy (Bag *et al*., 2015; Chen *et al*., 2009). Addition of histidine to patterned cells resulted in rapid and complete disassembly of the receptor clusters, demonstrating the reversibility of the systems, a key advantage of the approach to investigate receptor dynamics (**Figure 3C**).

**Figure 3.**
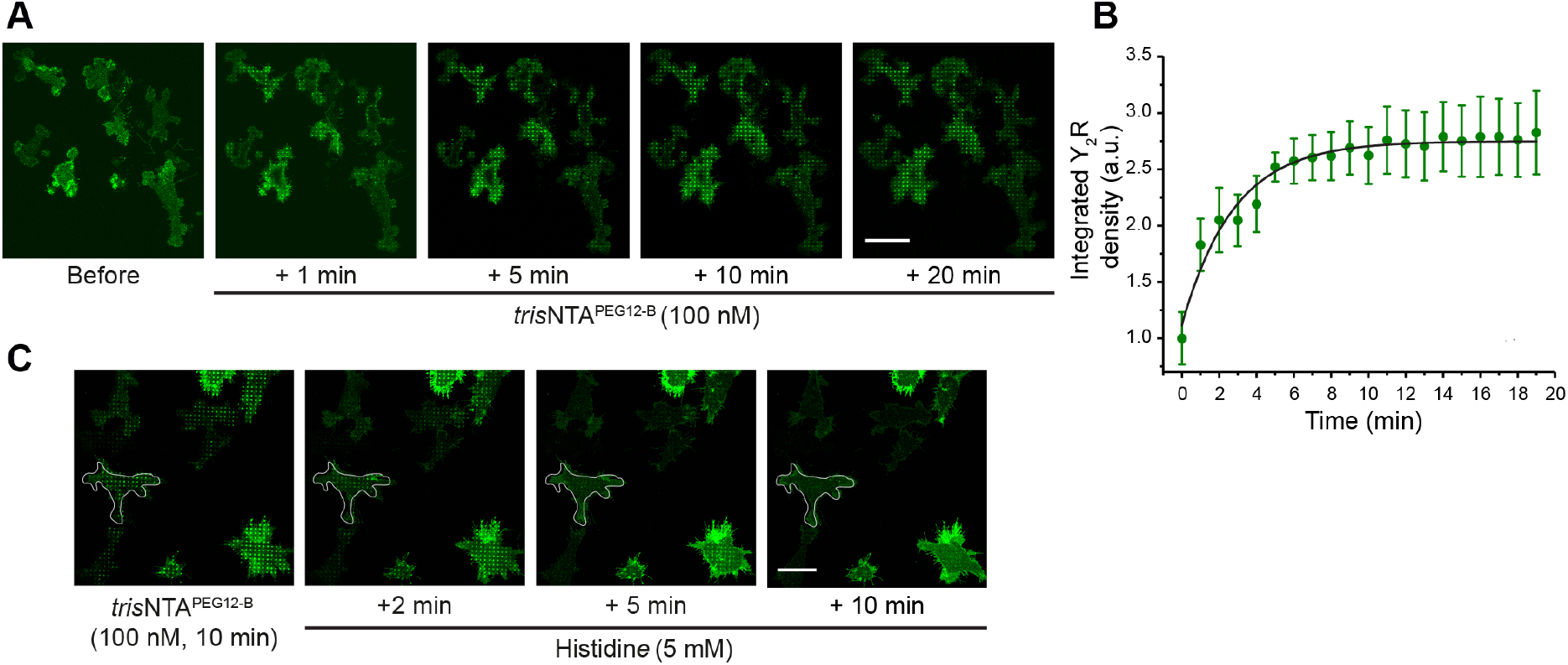
*In situ* receptor clustering with high spatiotemporal resolution. (A) Time-lapse imaging of Y_2_R assembly. Y_2_R-expressing HeLa cells were allowed to adhere to pre-structured SA-matrices for 3 h and were visualized by CLSM in LCIS at 37 °C. Time-lapse images were recorded for 20 min immediately after addition of *tris*NTA^PEG12-B^ (100 nM). Scale bar: 20 μm. (B) Receptor-integrated density in the patterned regions increased mono-exponentially, leading to an assembly rate of 0.35 ± 0.05 min^-1^ and τ_1/2_= 3 min. (50-200x 1 μm ROIs per experiment were analyzed from a total of 30 cells from three different experiments, 10 cells per experiment). (C) Reversal of the interaction and disassembly of the clusters is demonstrated upon addition of histidine. Y_2_R-expressing cells were allowed to adhere to the SA-matrices for 3 h, and then receptor confinement was induced by addition of *tris*NTA^PEG12-B^ (100 nM). Subsequently, cells were incubated with histidine (5 mM) for 2 to 10 min followed by washing. Scale bar: 10 μm.

Y_2_R activation by its natural ligand NPY promotes cell migration and proliferation (Ekstrand *et al*., 2003; Movafagh *et al*., 2006). In cells cultured on SA matrices, a 17% increase in cell area was detected after addition of the agonist porcine neuropeptide Y (pNPY, *K*_D_ = 5.2 ± 2.0 nM) (**Figure 4A, B**). When clustering was induced by the nanotool, we also observed a fast change in cell spreading and motility and a 20% increase in the total cell area concomitant to receptor assembly (**Figure 4A, C**). This analogous effect indicates a ligand-independent response to receptor clustering. We did not observe change in cell motility upon addition of the *tris*NTA^PEG12-B^ in cells cultured on matrices without SA. Furthermore, cells expressing Y_2_R^mEGFP^ (lacking a His_6_-tag) on SA-matrices showed no significant change in cell spreading upon addition of the nanotool, demonstrating the specificity of the response. (**Figure 4–figure supplement 1**). To investigate the relevance of clustering for Y_2_R activation and the cell motility response, we evaluated the increase in cell area upon ligand-induced activation in cells that were non-and pre-clustered by the nanotool. We revealed that nanotool-induced clustering amplified the motility effect induced by the pNPY ligand. In pre-clustered cells, stimulation with pNPY (10 nM) led to a 2-fold amplification and a 40% increase in cell area compared to the initial state (**Figure 4A, C**). A dose-dependent increase in cell area (**Figure 4A, C**) and cluster intensity (**Figure 4A, D**) was observed for *tris*NTA^PEG12-B^-pre-clustered cells. Overall, these results indicate a critical function of the receptor clusters, an amplification of the signal in pre-patterned cells, or, from the other point of view, a sensitization of the receptor to lower concentrations of the natural ligand NPY.

**Figure 4.**
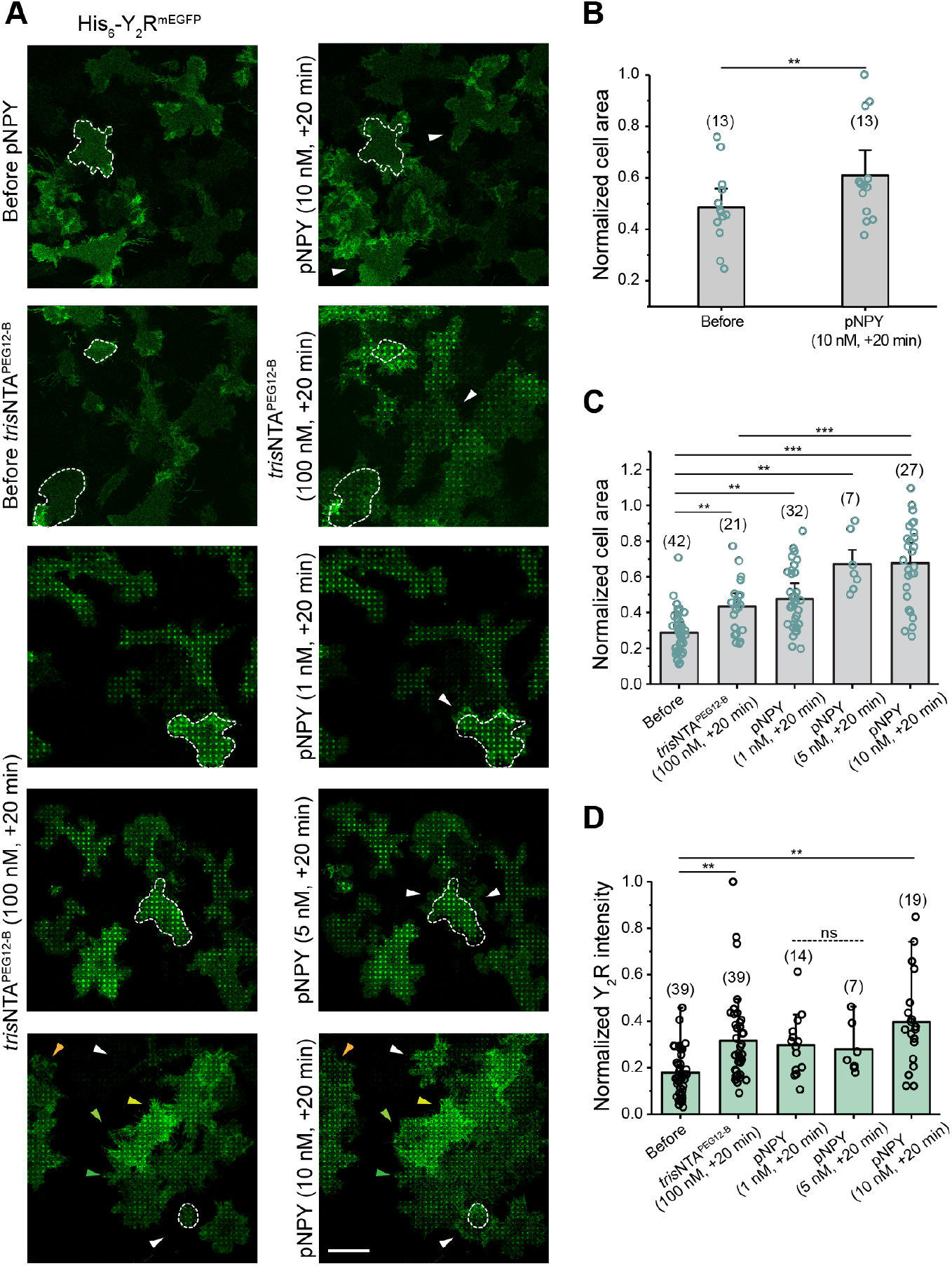
Receptor clustering amplifies the cell response induced by ligand activation. (A) Confocal microscopy images of cells expressing Y_2_R exposed to different conditions. Y_2_R-expressing HeLa cells were allowed to adhere to pre-structured SA-matrices for 3 h and visualized by CLSM in LCIS at 37 °C. Cells were visualized and imaged for 20 min after addition of *tris*NTA^PEG12-B^ or pNPY or both, first *tris*NTA^PEG12-B^ and subsequently pNPY (20 min incubation time, each). Scale bar: 20 μm. (B) Cell area analysis before and 20 min after addition of pNPY (10 nM) showed a 20% area increase, confirming an effect of ligand activation on cell motility. Values for cell area were normalized with respect to the highest value. The mean ± SD (13 cells) is shown. **p≤ 0.01 for Tukey test. (C) Cell area analysis before and 20 min after addition of *tris*NTA^PEG12-B^ (100 nM) and subsequent addition of pNPY (1, 5, and 10 nM, one well for each concentration) showed a dose-dependent area increase, demonstrating an amplification effect of receptor clustering in combination with pNPY. Values for cell area were normalized with respect to the highest value. The mean ± SD (42 cells before, 21 cells after *tris*NTA^PEG12-B^ and 14, 7, 19 for pNPY 1, 5, and 10 nM respectively) is shown. **p≤ 0.01 and ***p≤ 0.001 for Tukey test. (D) Quantification of receptor intensity in the nanotool-induced patterned regions showed a significant increase in pattern intensity after addition of pNPY (10 nM), the concentration that had the largest effect on cell motility. The mean ± SD is shown (19 to 39 cells and 50-220x 1 μm ROI, were analyzed). ***p≤ 0.001 for Tukey test.

As calcium signals are widely known to regulate cell motility, we monitored local calcium dynamics utilizing a far-red cell-permeable calcium-sensitive dye. By dual-color imaging, receptor assembly and the cytosolic calcium concentration were simultaneously recorded in living cells over the matrices. Upon addition of *tris*NTA^PEG12-B^, receptor recruitment led to a 2-fold increase in cytosolic calcium concentration with a rapid rise within two minutes (**Figure 5A, B**). A second peak in the cytosolic Ca^2+^ signals was detected upon subsequent addition of pNPY (10 nM). Contrary, no calcium signal was measured in cells over control matrices without SA (**Figure 5–figure supplement 1**). To confirm an enhancement of the response to ligand-induced activation in the presence of nanotool-induced receptor clusters, calcium signals were monitored in non-or pre-clustered cells (**Figure 5–figure supplement 2**). After receptor clustering, we observed a 1.6-fold increase in cytosolic calcium signal upon pNPY stimulation compared to the initial state. In contrast, a 1.2-fold increase was detected for cells in the presence of the pNPY only.

**Figure 5.**
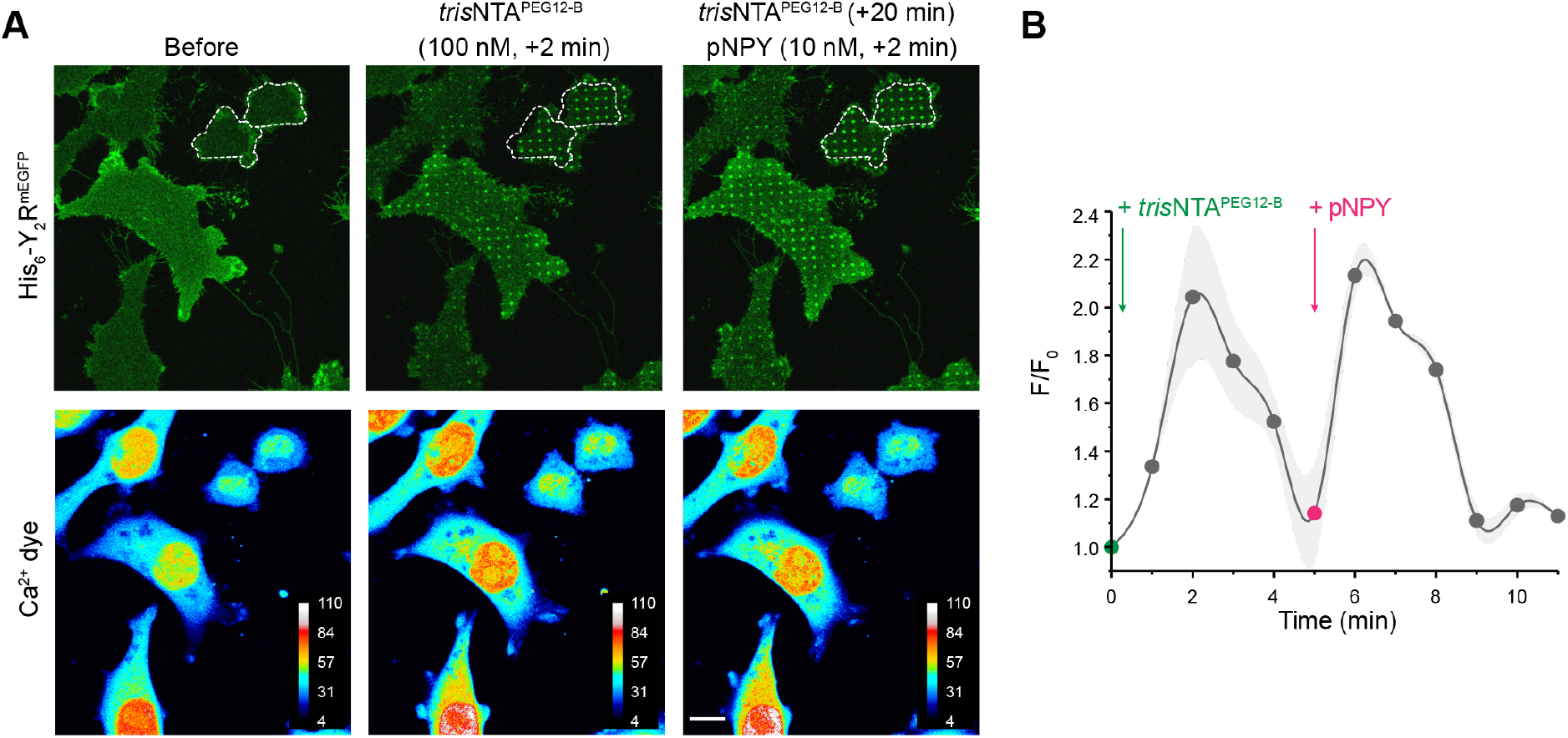
Ligand-free receptor confinement provokes calcium signaling. (A) Representative confocal fluorescence images of the Y_2_R (upper panel) and color-coded images of the Ca^2+^ dye (lower panel). Y_2_R-expressing cells on SA-pre-structured matrices were incubated with BioTracker 609 Red Ca^2+^ AM dye (3 μM) for 30 min. After rinsing, cells were immediately imaged by CLSM in LCIS at 37 °C. Addition of *tris*NTA^PEG12-B^ showed a 2-fold increase in cytosolic calcium. Scale bar: 10 μm. (B) Analysis of the mean gray value for Ca^2+^ signal before (F_0_) and upon addition of *tris*NTA^PEG12-B^ (F) *versus* time. Time-lapse images were recorded with 45 s interval before and after addition of *tris*NTA^PEG12-B^ (100 nM), and subsequent addition of pNPY (10 nM) after 5 min of nanotool addition (5 slices z-stack per time-point). ROIs covering the complete cell area were considered. The mean ± SD (10 cells) is shown.

Overall, our results show analogous calcium signaling for ligand-free *versus* ligand-induced systems and an amplification of the signal for ligand-induced activation in pre-clustered cells. Y_2_R has been found in a conformational equilibrium between inactive and active states in the absence of the ligand and forms high-affinity active complexes with G proteins (Ziffert *et al*., 2020). By ligand-free receptor clustering, the high local receptor density may increase the residence time of G proteins in vicinity and recruit further downstream effectors, which could boost the probability of activation and subsequent signaling. Based on the formation of the high affinity Y_2_/G protein complexes and the short time regime (1-5 min) in which changes in Ca^2+^ concentration and cell motility are observed, it is likely that the ligand-independent activation mechanism involves the G protein pathway. G protein signaling leads to the release of Gβγ and activation of phospholipase C-beta that cleaves phosphatidylinositol 4,5-bisphosphate into diacylglycerol and phosphatidylinositol (3,4,5)-trisphosphate (PIP_3_). PIP_3_ opens intracellular calcium stores through PIP_3_ receptors, leading to local activation of cytoskeletal proteins and causing the observed cell motility response.

### Ligand-free vs ligand-induced receptor activation differs in arrestin recruitment

We finally explored the impact of receptor clustering on downstream signaling by monitoring arrestin-3 recruitment. GPCR desensitization involves a complex series of events, e.g. receptor phosphorylation, arrestin-mediated internalization, receptor recycling, and lysosomal degradation (Ziffert *et al*., 2020). Short-term desensitization occurs within minutes and is primarily associated with arrestin preventing G protein interaction with the GPCR. Arrestins bind to activated, phosphorylated GPCRs and block receptor-G protein interaction by steric hindrance at the receptor-coupling interface, while serving as adaptors for key components of the endocytic machinery and numerous signaling proteins (Hilger *et al*., 2018; Wang *et al*., 2020). In the presence of high concentrations of the canonical ligand, an Arr3-dependent internalization, subsequent endosomal sorting, and recycling of Y_2_R to the cell membrane were observed (Walther *et al*., 2010; Wanka *et al*., 2018). However, recent studies demonstrated a strong and persistent activation of the G_αi_-pathway upon Y_2_R activation, which depletes the intracellular G protein repertoire before Arr3 binding can terminate signaling (Ziffert *et al*., 2020). To assess whether ligand-free clustering leads to Arr3 recruitment, we transfected cells stably expressing the Y_2_R with Arr3^mCherry^ (in brief Arr3) and monitored Arr3 recruitment in real-time by total internal reflection fluorescence (TIRF) microscopy (**Figure 6A**).

**Figure 6.**
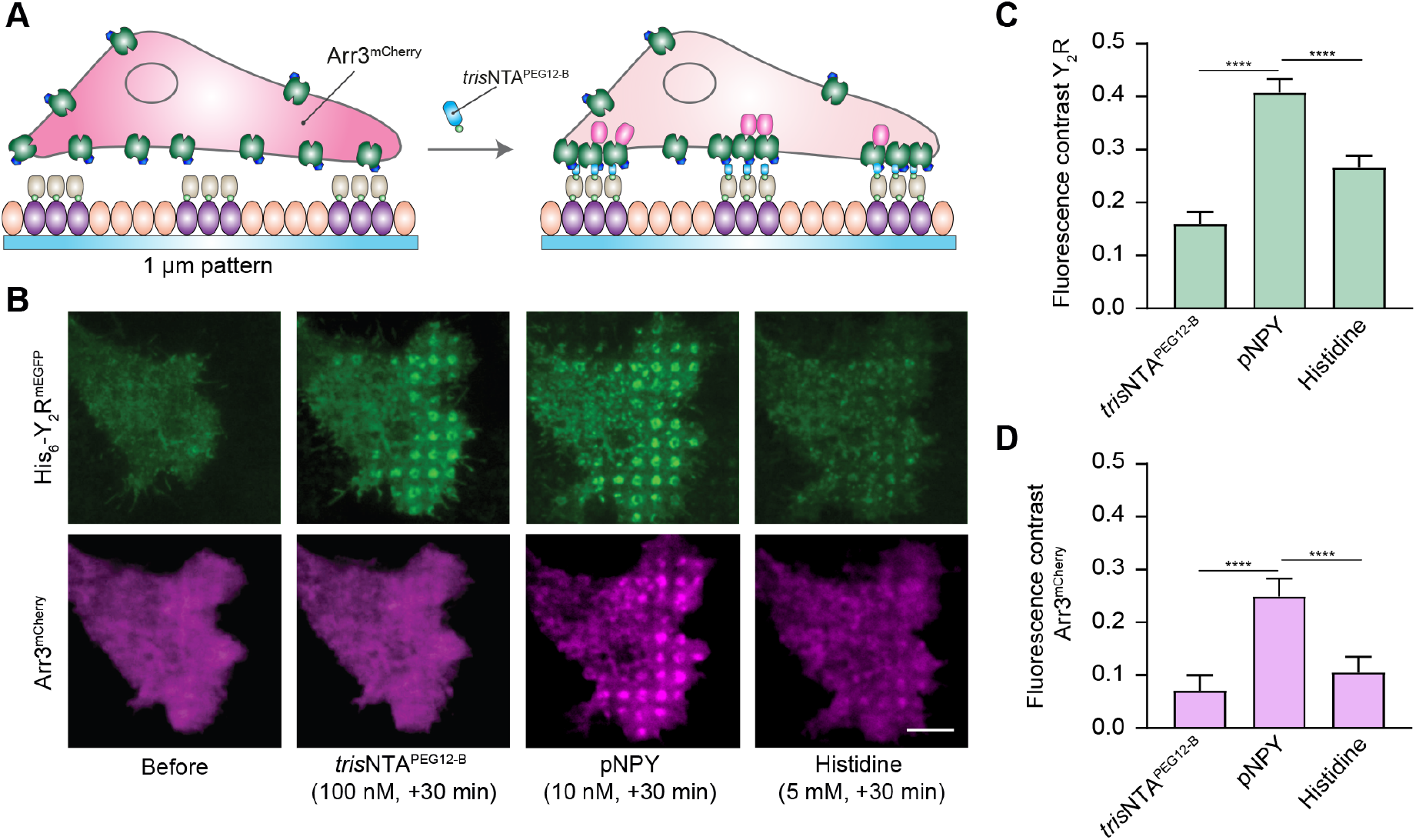
Arrestin-3 recruitment upon ligand-induced receptor activation. (A) Schematic representation of the experimental set-up. Cells co-expressing Y_2_R and Arr3 were allowed to adhere to SA-pre-structured matrices for 3 h and visualized by total internal reflection fluorescence (TIRF) microscopy in LCIS at 37 °C. (B) Representative TIRF images of cells before and upon addition of *tris*NTA^PEG12-B^ (100 nM, 30 min) and subsequent incubation with pNPY (10 nM) and histidine (5 mM) in LCIS for 30 min at 37 °C. All concentrations mentioned are final concentrations in the wells. Scale bar: 5 μm. (C) Quantification of the fluorescence contrast in the patterned regions for Y_2_R confirmed receptor enrichment upon addition of *tris*NTA^PEG12-B^, (2-fold with respect to the basal signal before, 100 nM, 30 min), which further increased 4-fold upon addition of pNPY (10 nM, 30 min). Histidine addition led to a decrease in the signal (1.7-fold decrease compared to pNPY, 5 mM, 30 min). Data were normalized with respect to the fluorescence intensity before clustering and it is expressed as the means ± SEM (60 cells for each condition were analyzed). Tukey’s multiple comparison test was applied (***p≤ 0.001). (D) Fluorescence contrast analysis demonstrated no significant recruitment of Arr3 upon *tris*NTA^PEG12-B^ (1.4-fold with respect to the basal signal before, 100 nM, 30 min). Addition of pNPY increased the Arr3 signal (3.6-fold, 10 nM, 30 min), confirming co-patterning of the downstream signaling molecules. Subsequent addition of histidine led to a decrease in the signal (2.3-fold, 5 mM, 30 min). Data was normalized with respect to the fluorescence intensity before clustering and it is expressed as the means ± SEM (60 cells for each condition were analyzed). Tukey’s multiple comparison test was applied (***p≤ 0.001).

In agreement with our results shown above, image analysis at an equilibrium state (30 min after addition of the nanotool) showed a subsequent increase in Y_2_R density in the clustered regions upon addition of pNPY (**Figure 6B, C**). Surprisingly, upon microscale receptor confinement by *tris*NTA^PEG12-B^, we did not observe a significant increase in Arr3 recruitment by intensity-contrast analysis of the patterned spots, whereas a significant Arr3 recruitment was detected upon addition of the agonist pNPY (**Figure 6B, D**). Reversibility by specific competition with histidine showed that half of the intensity in the patterned regions was dissipated of the Y_2_R/Arr3 assemblies (**Figure 6B, D**). These results suggest that not all receptors within the cluster regions are associated with the nanotool upon addition of the ligand, supporting the observation of increased receptor density in the presence of the pNPY. Patterning of Arr3 was also detected in cells on an anti-His_6_ antibody matrix within the first minutes after addition of pNPY (**Figure 6–figure supplement 1**). In this case, we did not observe a significant change in receptor density upon addition of the pNPY, indicating that the high degree of immobilization and large size of the antibody might restrict the transient enrichment of active receptors into the clustered regions. Specific clusters termed GPCR hot spots (40-300 nm) have been visualized at the plasma membrane of living cells (Calebiro & Jobin, 2019; Chavez-Abiega *et al*., 2019; Hilger *et al*., 2018; Sungkaworn *et al*., 2017). These hot spots represent regions that preferentially engage signaling, and that are enriched in both receptors and G proteins. We hypothesize that the induced microscale clusters trigger the formation of hot spots, which provide an ideal environment for recruitment of more active receptors and thus amplification of the signal. By increasing the local effective receptor concentration, this organization may amplify both the speed and efficiency of receptor-G protein coupling while enabling local signal transduction. In summary, our results show a difference between Arr3 recruitment in the ligand-free mode compared to the ligand-activated state. These observations indirectly confirm a high-affinity interaction between the Y_2_R and Gα_i_ and suggest active recruitment of G proteins that delay Arr3 recruitment and impair termination of G protein signaling (Ziffert *et al*., 2020). Likewise, the increased recruitment of receptors observed after addition of the pNPY ligand may be directly related to the dynamic nature of the confined regions.

## Discussion

We developed a versatile approach to cluster receptors *in situ* with minimal steric hindrance and disturbance. The transient association between the multivalent nanotool and the receptors revealed the generation of a dynamic platform for cell signaling. The dynamic exchange of molecules within induced Y_2_R microscale clusters may contribute to the formation of hot spots and final downstream signaling. This feature, as well as the broad applicability and the lower concentration required compared to established systems with immobilized ligands or antibodies, highlight the advantages of this versatile approach. Ligand-independent receptor activation by confinement was unraveled by cytosolic calcium increase and changes in cell spreading and motility, a response analogous to ligand-induced receptor activation. Furthermore, we demonstrated an amplification of the signal upon ligand-induced activation in cells pre-clustered with the nanotool. Subsequent addition of the neuropeptide ligand led to an enhancement of the calcium signal compared to ligand-induced activation without clustering. Interestingly, we demonstrated an increase in the receptor intensity in the clustered areas concomitant with ligand addition. We also uncovered a difference in downstream signaling for the ligand-free *versus* ligand-activated receptors as evidenced by co-recruitment of Arr3 to the clustered spots only occurring in the presence of the neuropeptide ligand. This finding is consistent with previous results demonstrating an Arr3-dependent internalization, subsequent endosomal sorting, and receptor recycling to the cell membrane in the presence of high concentrations of NPY (Walther *et al*., 2010; Wanka *et al*., 2018). We hypothesize that high-affinity Y_2_R/Gα_i_ interactions drive the initial cell response, cytosolic calcium increase, and cell motility. High local receptor density in the spots increases the residence time of proximate Gα_i_ proteins and recruits further downstream effectors, which boost the probability of activation (Sánchez *et al*., 2021). Further, Y_2_R/Gα_i_ interactions lead to persistent activation of the Gα_i_ pathway, which depletes the intracellular Gα_i_ protein repertoire before Arr3 binding can terminate signaling (Ziffert *et al*., 2020). The time frame of imaging after addition of the nanotool (30 min) suggests a long-lasting Gα_i_ protein activation and favors the hypothesis of a mechanism for Y_2_R activation and desensitization that is limited to the cell membrane and partially independent of Arr3 recruitment (Ziffert *et al*., 2020).

Multiscale analyses of the interactions between receptor clusters, G proteins, the lipid environment and actin-myosin assemblies are critical to confirm cluster behavior and dynamics. *In vitro* reconstitution systems utilizing lipid bilayers have proven useful to investigate receptor signaling (Huang *et al*., 2021). Comparing fluid-patterned lipid bilayers with our established platform together with advanced quantitative fluorescence microscopy techniques such as fluorescence resonance energy transfer (FRET) and single-molecule localization microscopy will help us to decode cluster behavior and decipher the complete ligand-independent signaling pathway. In summary, the developed nanotool and matrices allow the investigation of ligand-independent receptor activation *in situ*, facilitating the investigation of early key processes in cell signaling.

## Materials and Methods

### Synthesis of *tris*NTA^PEG12-B^

Cyclam-Lys-*tris*NTA (Gatterdam *et al*., 2018) (5.0 mg, 4.8 μmol), Biotin-PEG_12_-NHS (23.0 mg, 24.0 μmol) and DIPEA (12.2 μL, 72.0 μmol) were dissolved in0.5 ml dry DMF and stirred for 2 h at RT. After reaction, the volatile components were removed by lyophilization. Raw product was purified by reverse-phase (RP)-HPLC (mobile phase A: H_2_O + 0.1% TFA, B: CAN + 0.1% TFA; gradient 5% to 80% B in 20 min; MZ-PerfectSil, 300 ODS, 5 μm, 250 × 10 mm, flow 4 ml/min). A biotin moiety was integrated into the nanotool for immobilization to SA in the pre-structured matrices. The PEG_12_ linker between the biotin and the *tris*NTA unit increased the flexibility of the molecule. The identity of *tris*NTA^PEG12-B^ was confirmed by liquid chromatography-coupled mass spectrometry (LC-MS, Waters BioAccord System). Datasets were recorded with an ACQUITY UPLC I-Class Plus chromatography system and ACQUITY RDa Detector, which was set to a cone voltage of 25 V, capillary voltage of 1.2 kV and a desolvation temperature of 500 °C operating in positive ionization mode. For reverse-phase separation, an ACQUITY UPLC Peptide BEH C18 column (300 Å, 1.7 μm, 2.1 mm x 100 mm) was used (**Figure 1–figure supplement 4**). *tris*NTA^PEG12-B^ was dissolved in HBS buffer (20 mM HEPES-NaOH pH 7.5, 150 mM NaCl) and incubated with 10-fold excess of NiCl_2_. After 30 min incubation at 4 °C, the excess of Ni(II) was separated by a size exclusion chromatography gravity column (PD MidiTrap G-10).

### Microcontact printing

Large-area microcontact printing was performed as described previously (Lanzerstorfer *et al*., 2020) with modifications. In short, a field of a large-area PFPE elastomeric stamp (1 μm grid size), obtained by the EV-Group (St. Florian am Inn, Upper Austria, Austria), was cut out, and washed by flushing with ethanol (100%) and distilled water. After drying with nitrogen, the stamp was incubated in 50 ml BSA solution (1 mg/ml, Sigma-Aldrich) for 30 min followed by washing the stamp with phosphate-buffered saline (PBS) and distilled water. After drying with nitrogen, the stamp was placed with homogeneous pressure onto a clean epoxy-coated glass substrate (Schott Nexterion Slide E) and incubated overnight at 4 °C. The next day, the stamp was stripped from the glass with a forceps, and the microstructured glass was bonded to a 96-well plastic casting using an adhesive tape (3M) and closed with an appropriate lid.

### Functionalization of the pre-structured matrices

BSA-pre-structured wells were incubated with biotin-BSA (0.1 mg/ml, Thermo Fisher Scientific, Waltham, MA, USA) and SA (1 μM, Sigma-Aldrich, Munich, Germany) in PBS, each for 1 h at RT. Incubated wells were washed thoroughly with PBS after each step to remove unbound biotin-BSA and SA. For binding of soluble His-tagged proteins, wells were incubated with *tris*NTA^PEG12-B^ (0.5 μM) in HBS buffer for 1 h at RT. For nickel-loading, the pre-structured matrices were sequentially incubated with imidazole (1 M, 2 min), EDTA (100 mM, 2 min), and NiCl_2_ (10 mM, 5 min). Wells were carefully washed after each step. Finally, HBS buffer containing EDTA (50 μM) was used to remove the free, non-complexed nickel ions. His_6_-GFP (100 nM) previously expressed and purified was added to the wells and incubated for 30 min at RT. Experiments were performed in biological replicas (N=5).

### Cell culture

HeLa Flp-In™ T-Rex™ Y_2_R cells (His_6_-Y_2_R^mEGFP^ or Y_2_R^mEGFP^) were generated and cultured at 37 °C, 5% CO_2_, and 95% humidity (Sánchez *et al*., 2021). For culturing the stable cell line, high glucose Dulbecco’s modified Eagle’s medium (DMEM) (Gibco/Thermo Fisher Scientific) was supplemented with 10% tetracycline-free fetal calf serum (FCS, Bio&Sell), blasticidin S HCl (1 μg/ml, Thermo Fisher Scientific), and hygromycin B (50 μg/ml, Thermo Fisher Scientific). To induce receptor expression the cell medium was replaced with fresh medium containing tetracycline (0.1 μg/ml, Fluka) 18 h before imaging. The same concentration of tetracycline resulting in an efficient plasma membrane targeting was used for all the experiments. The cells were regularly tested for mycoplasma contamination.

### Receptor confinement in real-time by *tris*NTA^PEG12-B^

Cells expressing Y_2_R (His_6_-Y_2_R^mEGFP^ or Y_2_R^mEGFP^) were trypsinized and allowed to adhere to SA pre-structured matrices for 3 h or overnight. 15-18 h prior to the experiment, the cell medium was replaced with fresh medium containing tetracycline (0.1 μg/ml) to induce receptor expression. The cells were visualized by CLSM in live-cell imaging solution (LCIS, Thermo Fisher Scientific) at 37 °C. Cells were subsequently incubated with nickel-loaded *tris*NTA^PEG12-B^ (final concentration 100 nM) in LCIS for 10-15 min at 37 °C. Excess of unbound *tris*NTA^PEG12-B^ was removed by washing with LCIS. For reversibility experiments, micropatterned cells were incubated with histidine (5 mM) in LCIS for 2 to 10 min followed by washing with LCIS. Experiments were performed in biological replicas (N=4).

### Receptor confinement on antibody-micropatterned matrices

Wells pre-structured with BSA were subsequently incubated with biotin-BSA (0.1 mg/ml), SA (1 μM), and a biotinylated anti-His_6_ antibody (1 μM) (ab106261, Abcam) in PBS for 1 h at RT. Wells were washed thoroughly with PBS to remove unbound antibody. Cells expressing Y_2_R were trypsinized and seeded onto the antibody patterns. After 3 h, cells were visualized by CLSM in LCIS at 37 °C. Experiments were performed in biological replicas (N=5).

### Time-lapse calcium imaging

18 h after seeding the cells onto pre-structured SA-matrices, cells were incubated with BioTracker 609 Red Ca^2+^ AM dye (3 μM, Merck Millipore) in fresh medium for 30 min. The cell-membrane permeable dye is de-esterified by cellular esterases and remains trapped in the cytosol. After incubation with the Ca^2+^ dye, cells were rinsed three times with PBS and imaged by CLSM in LCIS at 37 °C. For investigation of Ca^2+^ signal, time-lapse images were taken (5 slices z-stacks, 45-s interval) before and after addition of *tris*NTA^PEG12-B^. Fluorescence intensity (λ_ex/em_ 590/609 nm) of the dye changes depending on the intracellular Ca^2+^ concentration. Maximum intensity projections of single channels were analyzed. The ImageJ ROI tool was used to define the areas of the image to be analyzed. We consider a ROI covering the complete cell contour. Mean gray values (F) were background subtracted and normalized to the fluorescence in cells before F_0_. Experiments were performed in biological replicas (N=3).

### Plasma membrane staining

Live-cell membrane staining was performed directly after receptor assembly in living cells grown on pre-structured matrices. CellMask™ deep red plasma membrane stain (Thermo Fisher Scientific) was used according to manufacturer’s instruction. 1 μl of the stock solution (1000x dilution) was dissolved in 1 ml of warm LCIS (final concentration 5 μg/ml) and subsequently added to the cells, incubated for 5 min at 37 °C, and washed with LCIS before visualization. Experiments were performed in biological replicas (N=3).

### Confocal laser scanning microscopy

Images were recorded by using a CLSM Zeiss LSM 880 (Carl Zeiss) equipped with a Plan-Apochromat 63x/1.4 Oil DIC M-27 objective. Sequential settings for dual-color imaging were used. Excitation wavelengths for the different fluorophores: 488 nm (argon laser) for mEGFP; 594 nm for the Ca^2+^ dye; 633 nm (helium-neon laser) for the plasma membrane dye. Signals were detected after appropriate filtering on a photomultiplier. Intensities of channels were adjusted over the whole image for better visualization of overlap and exported by Zen blue (version 2.3 lite, Zeiss). Detector amplification, laser power, and pinhole were kept constant for all studies.

### Image analysis

Fluorescence images were processed with Zen blue, ImageJ, and Fiji software (Schindelin *et al*., 2012; Schneider *et al*., 2012). All images were background subtracted. Integrated density, mean gray value and cell area were obtained with ImageJ. Data were plotted with OriginPro.

### Fluorescence recovery after photobleaching

FRAP experiments were conducted at the CLSM Zeiss LSM 880 using 63 x/1.4 Oil DIC objective. Rectangular-shaped regions (6-10 μm radius) were bleached within 10 s with high laser intensities. Fluorescence recovery was monitored by repetitively imaging an area containing the photobleached region at 0.1 frame/s for ∼150 s. For the analysis, a simulation approach that allows computation of diffusion coefficients regardless of bleaching geometry used in the FRAP series was applied (Blumenthal *et al*., 2015). The method is based on fitting a computer-simulated recovery to actual recovery data of a FRAP series. The algorithm accepts a multiple-frame TIFF file, representing the experiment as input, and simulates the diffusion of the fluorescent probes (*2D* random walk) starting with the first post-bleach frame of the actual data. Once the simulated recovery is finished, the algorithm fits the simulated data to the real one and extracts the diffusion coefficient. The algorithm iteratively creates a series of simulated images, where each frame corresponds to a single iteration. The intensity values are extracted from the (user indicated) bleached area of the simulated frames, thus determining the general shape of the recovery curve. The “time” axis at this stage is in arbitrary units (iterations). To extract the diffusion coefficient, the simulated recovery curve needs to be fitted to the real recovery curve, by appropriately stretching the “time” axis. The time between frames in the actual data set is obviously known, thus once overlapping optimally the simulated curve with the real one, the duration of one iteration, in real-time units, is determined. The diffusion coefficient of the simulated series is then calculated according to eq. 1, where *D*_*s*_ is the simulation-extracted diffusion coefficient, *l* is the step of a molecule in each iteration of the simulation, corresponding to one pixel in the image (the pixel size is calibrated previously, by imaging a known calibration sample), and *t*_*i*_ is the time interval between steps (determined as explained).

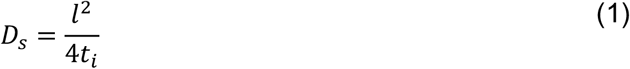

The simulation proceeds until a plateau is reached (equilibration of the fluorescence intensity in the bleached area). The number of data points in the simulated recovery is typically different (larger) than the number of experimental points. In addition, the real experimental data may not have been acquired until equilibration of fluorescence. To determine *t*_*i*_, the algorithm scans a range of possible values for the total duration represented by the simulation and calculates a value *X*^2^ for the goodness-of-fit between the simulated data and the real FRAP data. Total simulation duration is selected as the one that produces the minimal *X*^2^. Experiments were performed in biological replicas (N=3).

### imFCS analysis

imFCS measurements were performed as described earlier (Harwardt *et al*., 2018; Harwardt *et al*., 2017). A home-built widefield setup with total internal reflection fluorescence (TIRF) illumination was used for imFCS analysis. The experimental setup was equipped with a 488 nm diode laser (100 mW, Obis, Coherent, USA). The excitation light passes through an acousto-optical tunable filter (AA Opto-Electronic, Orsay, France) and a telescope consisting of two achromatic lenses (Thorlabs, USA) with f = −40 mm and 750 mm. A third achromatic lens (f = 400 mm, Thorlabs) directed the excitation light to the TIRF mirror and had its focus on the back focal plane of the objective. The TIRF mirror was placed on a motorized translation stage (25 mm, #MTS25/M-Z8, Thorlabs) controlled by a motion controller (K-Cube Brushed DC Servo Motor Controller, #KDC101, Thorlabs) to switch between widefield and TIRF illumination. The light entered an Eclipse Ti microscope (Nikon, Japan) was reflected by a dichroic mirror (TIRF-Quad filter set 405/488/561/640 consisting of a QuadLine Laser Clean-up ZET405/488/561/640x, QuadLine dichroic zt405/488/561/640rpc, QuadLine rejection band ZET405/488/561/640 TIRF, all AHF Analysentechnik AG, Tübingen, Germany), and was directed onto the sample by an oil-immersion TIRF objective (UapoN 100xOTIRF, 1.49 Oil, Olympus, Japan). A nosepiece stage (IX2-NPS, Olympus) was used for z-plane adjustment and drift minimization. Emission light was collected by the same objective and passed the dichroic mirror. In the detection path a TwinCam (Acal Bfi, Germany) with a BrightLine HC 525/45 bandpass filter (AHF Analysentechnik AG) was implemented, and the signal was detected by a scientific complementary metal-oxide semiconductor (sCMOS) camera (Zyla 4.2, Andor, Belfast, UK). Data were collected using the open-source software μManager (Edelstein *et al*., 2010). For data acquisition the following settings were applied: 24 W/cm^2^ laser intensity, a bit depth of 16 bit, pixel readout rate of 540 MHz, frame time 4 ms, 4×4 binning, and 4,000 frames per film. For each film, a 40×25 pixel (or 40×20 pixel) region of interest (ROI) was chosen and the measurement was performed with TIRF illumination to observe membrane diffusion of Y_2_R. In total 36 untreated cells, 24 cells with *tris*NTA^PEG12-B^ immobilized receptors, and 26 cells with anti-His_6_ antibody immobilized receptors were measured. Each condition contains data from at least three independent measurement days (N=3).

### imFCS data analysis

Analysis of imFCS films was performed using the imFCS plugin (version 1.52) (Sankaran *et al*., 2010) for Fiji (Schindelin *et al*., 2012). The following correlation settings were chosen: emission wavelength = 515 nm, NA = 1.49, correlator scheme P = 16 and Q = 8, lateral PSF = 0.8, binning = 1, pixel size = 5.75 μm, magnification = 25 for 4×4 binning, and linear segment bleach correction with linear segments of 500 frames. Diffusion coefficients were obtained for each pixel by fitting the correlation curves according to the literature (Sankaran *et al*., 2010). To compare the overall diffusion coefficients with those of the patterned regions, ROIs were placed around patterned regions and analyzed separately. For further analysis, the pixelwise diffusion coefficients for all measurements were imported into OriginPro 2019 (OriginLab Corporation, Northampton, USA). For box plots of diffusion coefficients, median diffusion coefficients were determined for each cell. Mean diffusion coefficients per condition were obtained by averaging over the median diffusion coefficients per measurement and calculating the standard error of the mean. Two-sample t-tests (α = 0.05) were applied to compare the diffusion coefficients for the different conditions. All datasets were tested for normality using the Kolmogorov-Smirnov test (α = 0.05). Significance was assigned as follows: p > 0.05 no significant difference between populations (n.s.), p < 0.05 significant difference (*), p < 0.01 significant difference (**), and p < 0.001 significant difference (***). Two-dimensional maps of diffusion coefficients were generated also in OriginPro. Diffusion coefficients were color-coded from light yellow to dark red in the range of 0 to 0.5 μm^2^/s. Pixels that yielded correlation curves with diffusion coefficients higher than 0.5 μm^2^/s are presented in black. Pixels that yielded correlation curves which could not be fitted by the imFCS plugin in Fiji are shown in light grey. To generate frequency distribution plots, diffusion coefficients were log-transformed and binned in the interval between −5.3 and 1.0 with a bin size of 0.1 for each cell. Logarithmic diffusion coefficients were re-transformed, frequency counts were averaged over all cells per condition, and normalized. Frequency counts were plotted logarithmically against diffusion coefficients. Errors bars represent standard errors of the mean.

### Arr3 recruitment upon receptor confinement

Microstructured surfaces were functionalized with biotin-BSA and SA or SA and anti-His_6_ antibody as described before. For transient co-transfection with Arr3^mCherry^, cells were sub-cultured the day before and then transfected with the Arr3^mCherry^ plasmid using the TurboFect™ transfection reagent (Thermo Fisher Scientific), according to the manufacturer’s instructions and induced with Tetracycline (0.1 μg/ml) 18 h before microscopy. Cells co-expressing His_6_-Y_2_R^mGFP^ and Arr3 were seeded onto the microstructured matrices and visualized by total internal reflection fluorescence (TIRF) microscopy in LCIS at 37 °C after 3 to 4 h to ensure a homogeneous cell membrane adhesion, which is a prerequisite for quantitative TIRF microscopy. For antibody experiments, cells grown on pre-structured matrices were incubated with pNPY (10 nM, Tocris) in LCIS for 30 min at 37 °C. For *tris*NTA^PEG12-B^ experiments, cells grown on SA-matrices were subsequently incubated with nickel-loaded *tris*NTA^PEG12-B^ (100 nM final) and pNPY (10 nM final) in LCIS for 30 min at 37 °C. For reversibility, cells were incubated with histidine (5 mM) in LCIS for 30 min. Experiments were performed in biological replicas (N=2).

### Arr3 imaging by TIRF microscopy

The detection system was set up on an epi-fluorescence microscope (Nikon Eclipse Ti2). For selective fluorescence excitation of mGFP and mCherry, a multi-laser engine (Toptica Photonics, Munich, Germany) was used at 488 and 561 nm, respectively. The samples were illuminated in total internal reflection (TIR) configuration (Nikon Ti-LAPP) using a 60x oil immersion objective (NA = 1.49, APON 60XO TIRF). After appropriate filtering using standard filter sets, the fluorescence was imaged onto a sCMOS camera (Zyla 4.2, Andor, Northern Ireland). The samples were mounted on an x-y-stage (CMR-STG-MHIX2-motorized table, Märzhäuser, Germany), and scanning was supported by a laser-guided automated Perfect Focus System (Nikon PFS).

### Contrast quantification and statistical analyses

Contrast analysis was performed as described previously (Lanzerstorfer *et al*., 2014; Lanzerstorfer *et al*., 2020; Schütz *et al*., 2017). Initial imaging recording was supported by the Nikon NIS Elements software. Images were exported as TIFF frames and fluorescence contrast analysis was performed using the Spotty framework (Borgmann *et al*., 2012). The fluorescence contrast <c> was calculated as <c> = (F^+^ - F^−^)/(F^+^ - F_bg_), where F^+^ denotes the intensity of the inner pixels of the pattern. F^−^ shows the intensity of the surrounding pixels of the micropattern, and F_bg_ the intensity of the global background. Data are expressed as the means ± SEM. Comparisons of more than two different groups were performed using one-way ANOVA, which was followed by Tukey’s multiple comparisons test in GraphPad Prism software (version 9.1.2).

## Acknowledgements

We thank Christian Winter, Andrea Pott, Inga Nold, and all members of the Institute of Biochemistry (Goethe University Frankfurt) for discussion and comments. We thank Dr. Annette Beck-Sickinger (Leipzig University) for the Y_2_ receptor construct and Dr. Alina Klein (Goethe University Frankfurt) for the generation of the Y_2_R^mEGFP^ constructs with and without His_6_-tag. We tank Dr. Cornelius Krasel (Philipps University of Marburg) for the Arr3^mCherry^ construct. We also thank Christian Winter for the LC-MS analysis. This work was supported by the German Research Foundation (GRK 1986 (No. 237922874) to R.W. and R.T.; CRC 807 (No. 57566863) P16 to R.T.), LOEWE DynaMem P3 to R.W. and R.T., the Volkswagen Foundation (Az. 96 498 to R.W., Az. 96 497 to M.H., and Az. 96 496 to R.T.); the Christian-Doppler Forschungsgesellschaft (Josef Ressel Centre for Phytogenic Drug Research, the Austrian Science Fund (FWF, I4972-B) and the FH Upper Austria Center of Excellence for Technological Innovation in Medicine (TIMed Center) to U.M., J.W. and P.L..

## Author contributions

M.F.S. performed the cell-based assays and imaging experiments. M.S.D. carried out the imFCS experiments and analyzed the data together with M.H.. U.M., P.L., and J.W. prepared the pre-structured surfaces, performed the Arr3 recruitment assays and the intensity-contrast analysis. K.G. synthesized and characterized the chelator compound. M.F.S., R.W., and R.T. wrote the manuscript with contributions from all authors. R.T. conceived the study.

## Competing interest

The authors declare no competing interest.

## Data availability

Data and movies are available in the supplementary materials.

## Supplemental figures

**Figure 1–figure supplement 1.**
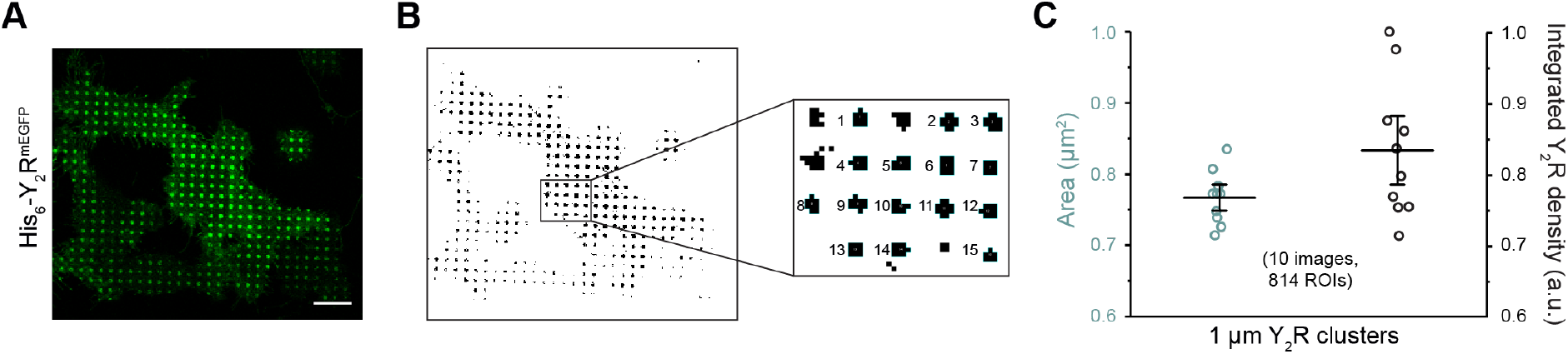
Receptor confinement with high reliability. (A) Representative confocal image of a cell patterned by *tris*NTA^PEG12-B^. (B) Automatic cluster analysis performed by ImageJ requires a “binary”, black and white, image. A threshold range is set to select the objects of interest apart from the background. All pixels in the image whose values lie under the threshold are converted to white and all pixels with values above the threshold are converted to black. Further selection of the clusters according to area and roundness enable a large-scale analysis. (C) Change in cluster area and integrated density of the receptor within different 96-well plates, different months and cell stocks reflected a reliable and reproducible approach. The average area in the clustered regions (0.77 ± 0.03 μm^2^) and integrated density of ten images in five different experiments (841x ROIs of 1 μm in total) is shown. Scale bar: 10 μm.

**Figure 1–figure supplement 2.**
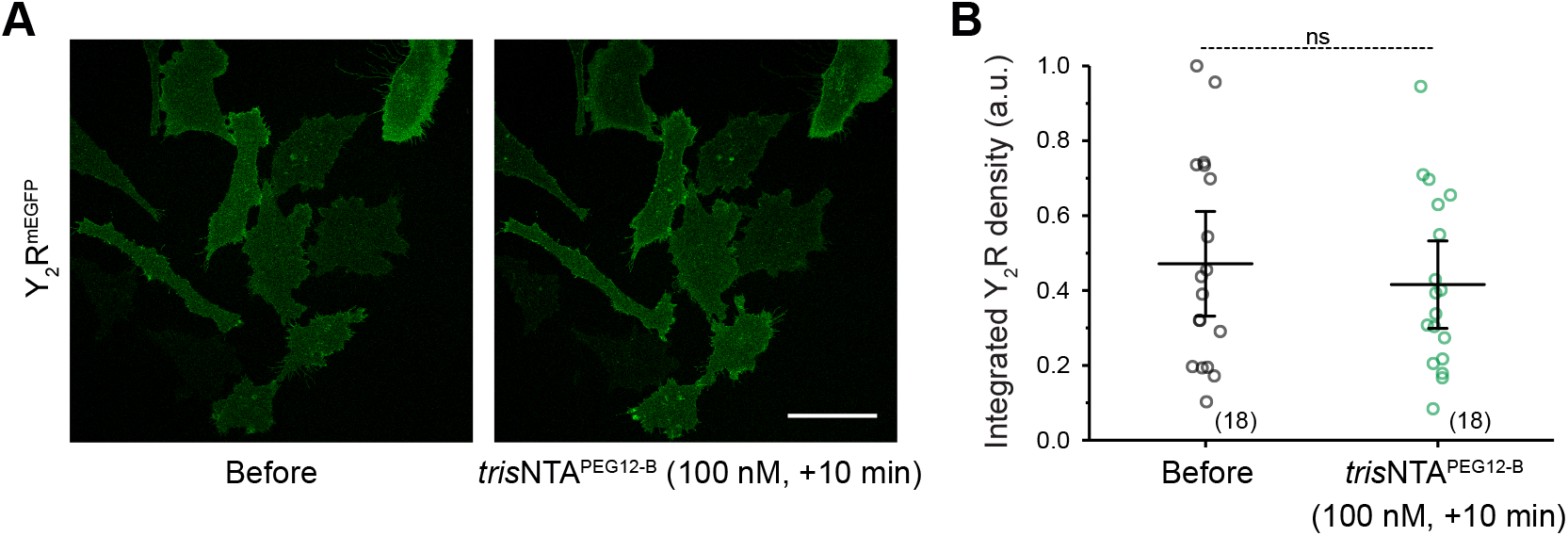
Y_2_ receptors lacking a His_6_-tag do not cluster in confined areas. (A) Representative confocal images of cells expressing Y_2_ receptors without N-terminal His_6_-tag over SA-pre-structured matrices before and after addition of the nanotool. Within the timeframe of imaging, there was neither a pattern formation nor a change in the integrated receptor density. (B) Quantification of the integrated Y_2_R density before and after addition of *tris*NTA^PEG12-B^. The mean ± SD (18 cells) is shown. **p≤ 0.01 for Tukey test. Scale bar: 50 μm.

**Figure 1–figure supplement 3.**
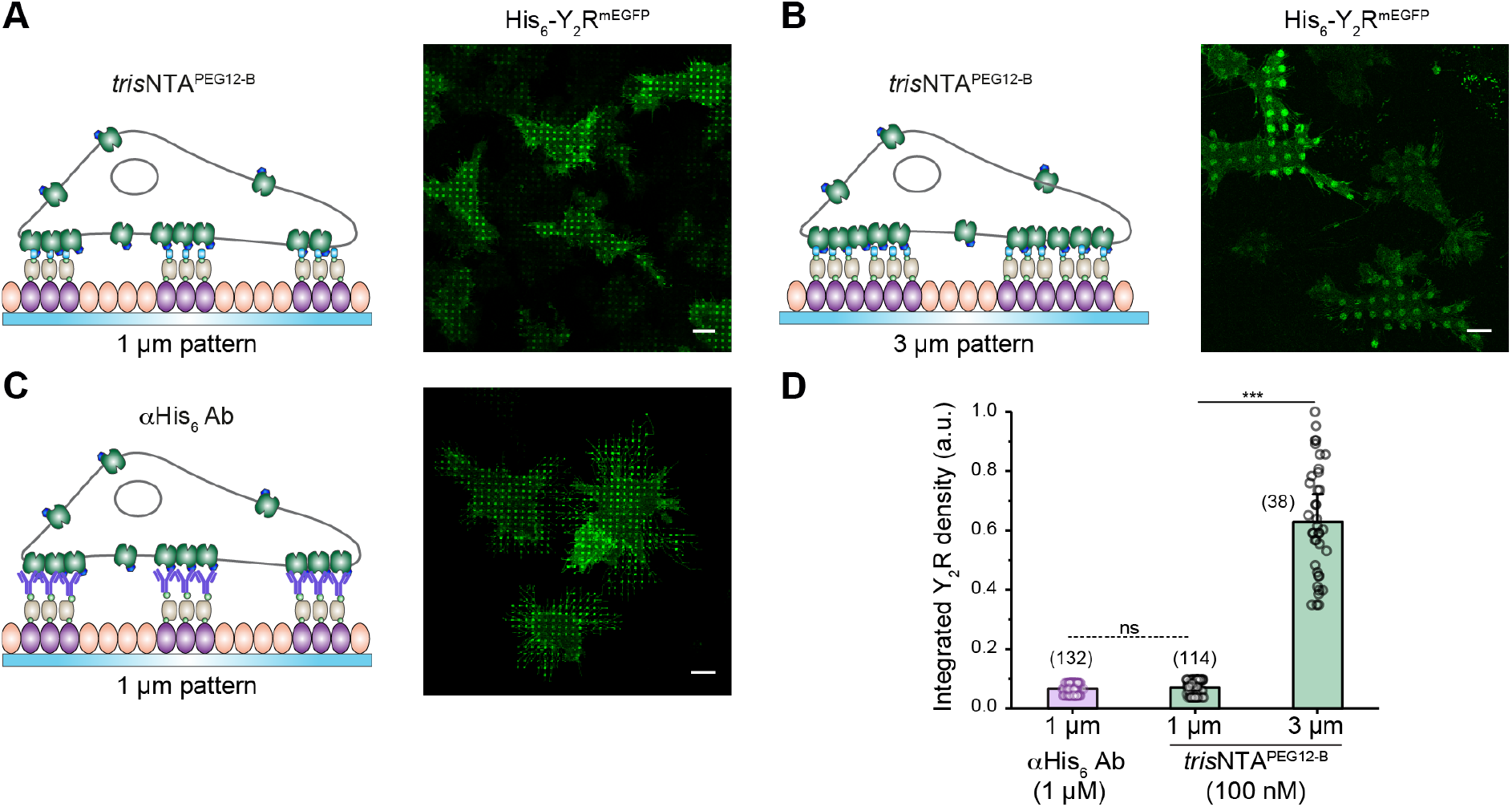
Receptor density correlates with the area of the pre-structured regions. (A, B) BSA-pre-structured matrices, 1 μm (A) or 3 μm (B), were stepwise functionalized with biotin-BSA and SA. Y_2_R-expressing HeLa cells were allowed to adhere to the functionalized matrix for 3 h and immediately imaged by CLSM in live-cell imaging solution (LCIS) at 37 °C. Incubation with *tris*NTA^PEG12-B^ (100 nM final) led to *in situ* receptor assembly. (C) 1 μm BSA-pre-structured matrices were stepwise functionalized with biotin-BSA, SA, and a biotinylated anti-His_6_ antibody. Y_2_R-expressing cells were allowed to adhere to the functionalized matrix for 3 h and immediately imaged by CLSM in LCIS at 37 °C. (D) Quantification of the receptor-integrated density. *In situ* receptor confinement by *tris*NTA^PEG12-B^ resulted in a receptor density that is comparable to cells in contact with pre-structured antibodies. For the 3 μm patterns, receptor density correlated with pattern area. The mean ± SD (38 to 132x 1 μm ROIs) is shown. ***p≤ 0.001 for Tukey test. Scale bars: 10 μm.

**Figure 1–figure supplement 4.**
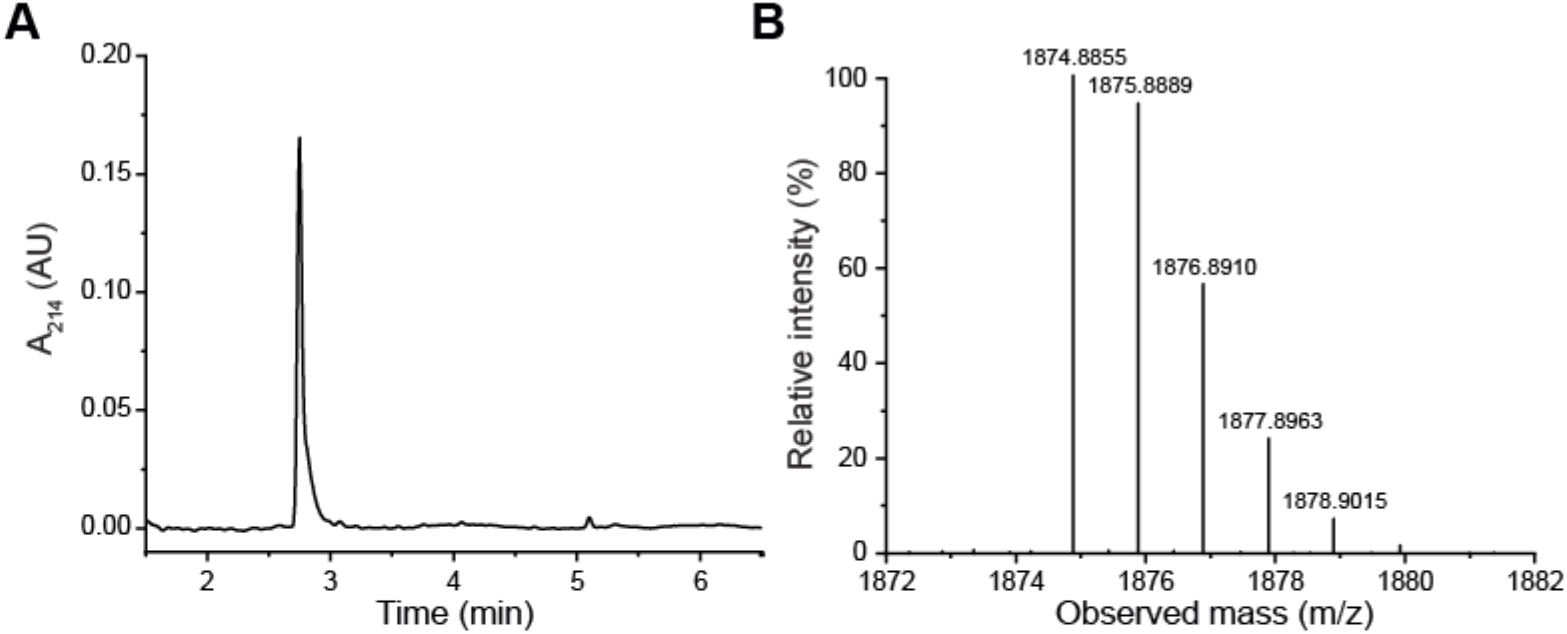
Multivalent nanotool *tris*NTA^PEG12-B^ analyzed by LC-MS. (A) *tris*NTA^PEG12-B^ chromatogram reflecting the purity of the synthesized nanotool. (B) LC-MS of *tris*NTA^PEG12-B^, yielding the experimental mass (M_exp._) of 1874.85 Da (M_theor._ = 1873.90 Da).

**Figure 2–figure supplement 1.**
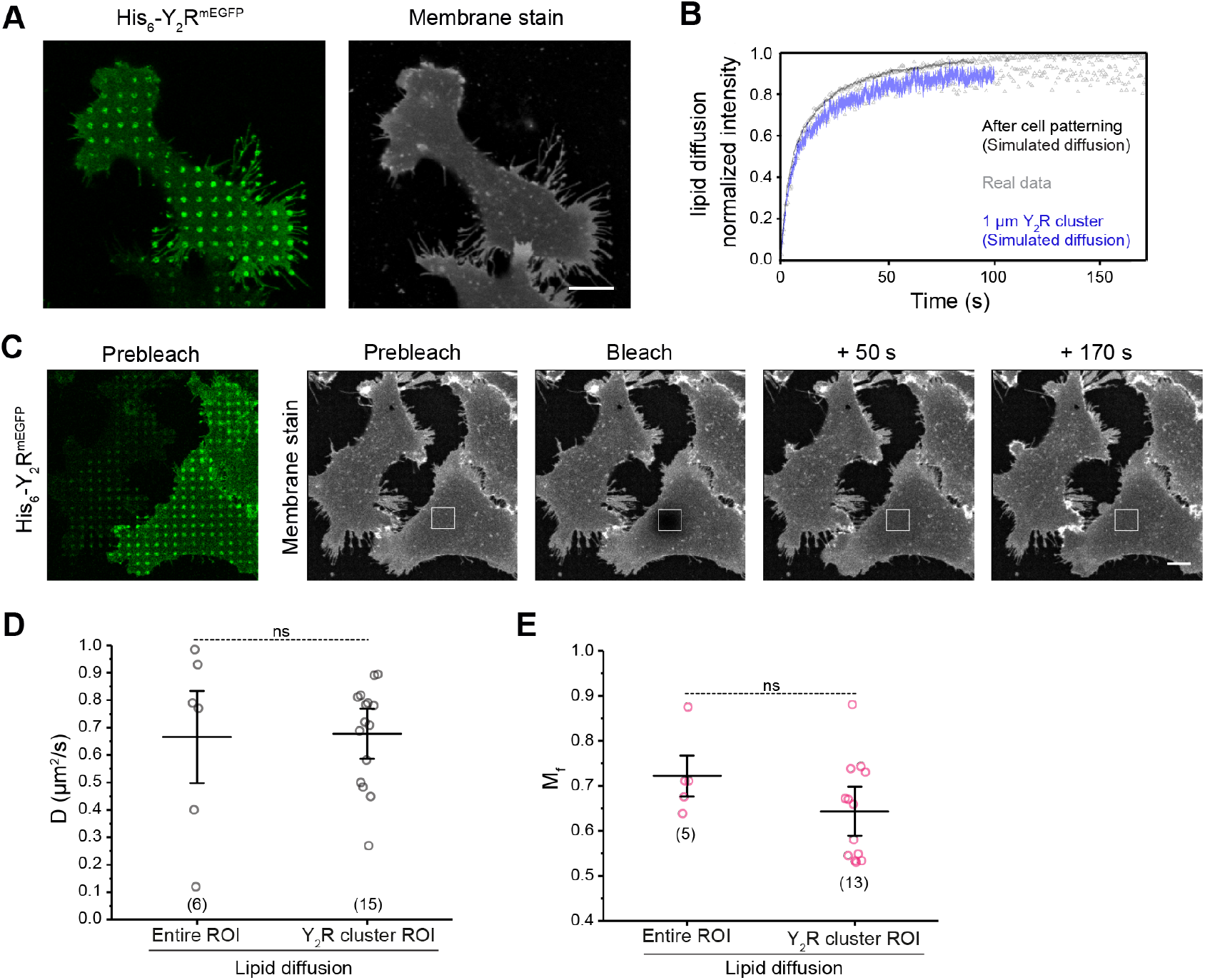
Lipid localization and dynamics after receptor confinement. (A) Confocal microscopy images of the live-cell plasma membrane staining, which was performed 15 min after Y_2_R assembly in living cells. 5 μg/ml CellMask staining solution was incubated for 5 min at 37 °C and washed with LCIS before visualization. Lipid distribution is not affected by receptor confinement as shown by the homogeneous staining of the membrane. (B, C) FRAP recovery curve (B) and time-lapse (C) for the lipid dye demonstrated a rapid recovery for the lipids. Diffusion was measured in the entire rectangular ROI or at the Y_2_R cluster spots (region selected based on the receptor channel image). An image of the receptor channel confirmed the presence of the pattern. (D) The analysis did not show any differences in lipid diffusion coefficients for the entire rectangular ROI or at the 1 μm clustered regions (*D*_entire ROI_ = 0.66 ± 0.10 μm^2^/s and *D*_spots_ = 0.67 ± 0.17 μm^2^/s). The mean ± SD (6 cells, 15x 1 μm ROIs) is shown. **p≤ 0.01 for Tukey test. (E) Quantification of the mobile fraction (*M*_f_) for FRAP measurements of the lipid dye reflected no significant difference. The mean ± SD (5 cells, 14x 1 μm ROIs) is shown. **p≤ 0.01 for Tukey test. Scale bars: 10 μm.

**Figure 2–figure supplement 2.**
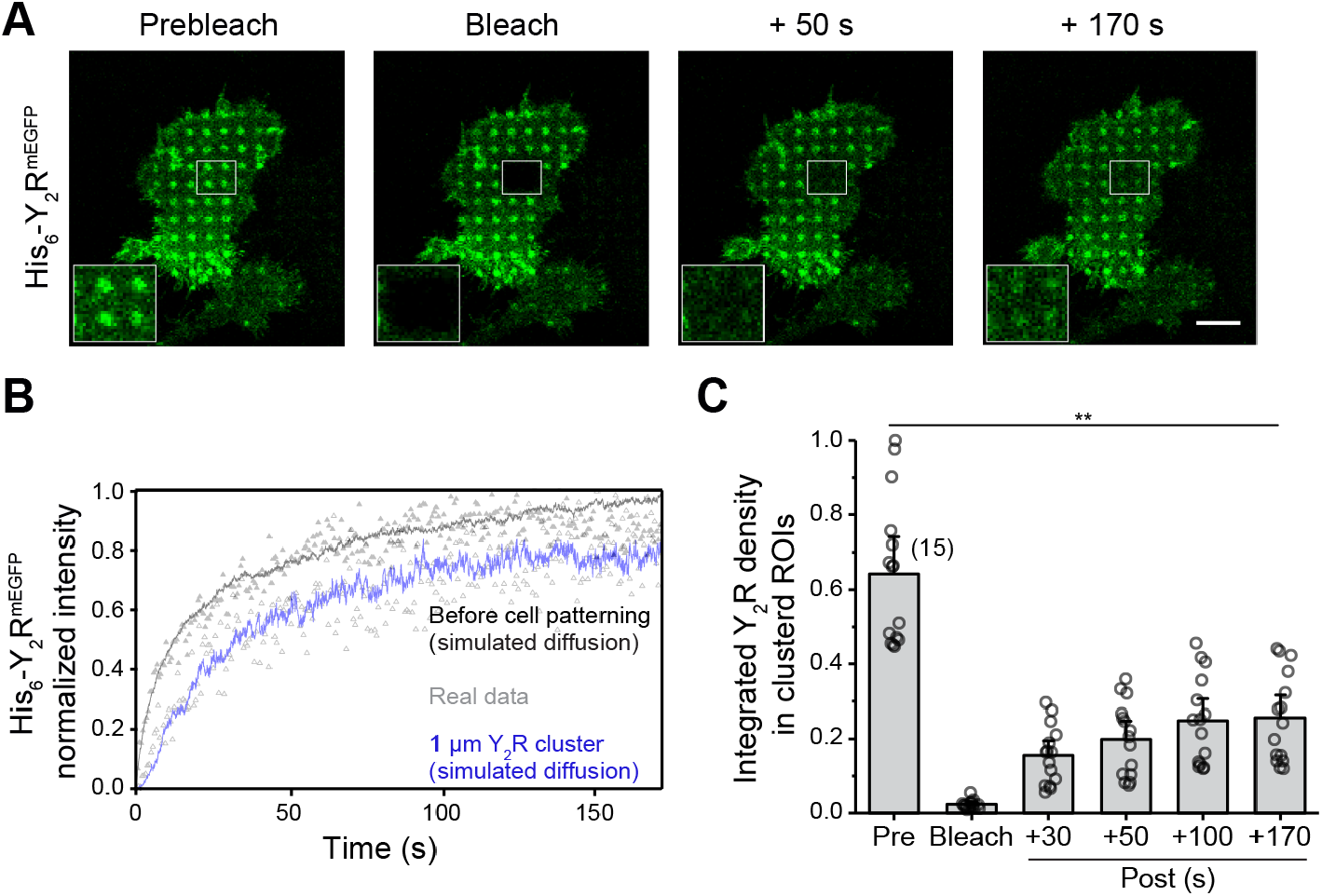
Dynamic receptor exchange in confined clusters. (A) Representative confocal images of FRAP measurements for Y_2_R-expressing cells on SA-pre-structured matrices 10 min after addition of the nanotool. (B) FRAP recovery curves reflecting the entire bleached area or an analysis performed only in the clustered 1 μm regions. The analysis is based on a simulation approach which fits a computer-simulated recovery to actual recovery data of a FRAP series and determines the diffusion coefficient regardless of bleaching geometry. (C) Quantification of the receptor density in the confined regions showed 50% recovery indicating a high exchange rate. The mean ± SD (6 cells, 15x 1 μm ROIs analyzed) is shown. **p≤ 0.01 for Tukey test. Scale bar: 10 μm.

**Figure 2–figure supplement 3.**
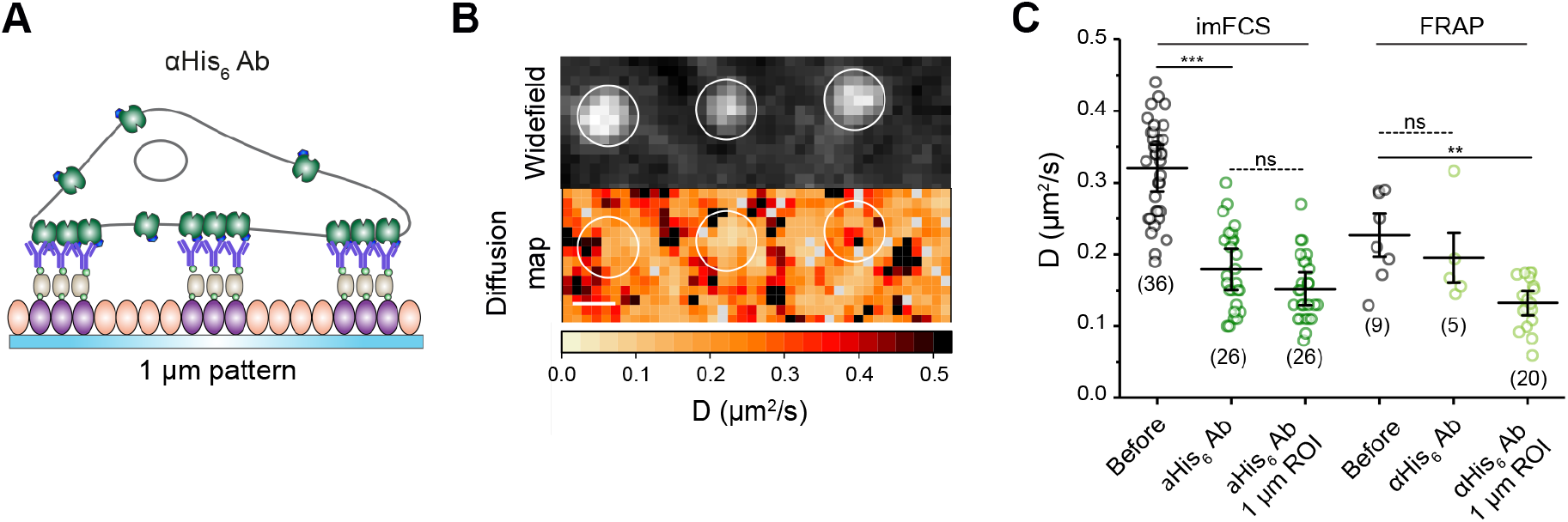
Receptor mobility on antibody structured matrices. (A) Scheme representing the experimental set-up. (B) Representative widefield image (left) of a ROI at the plasma membrane of a living cell over a pre-structured matrices with an anti-His_6_ antibody analyzed by imFCS and the derived two-dimensional diffusion map (right). (C) Lateral diffusion of the receptor analyzed by FRAP and imFCS. Both techniques demonstrated a decrease in D at the plasma membrane (*D*_*before*_= 0.32 ± 0.06 μm^2^/s and 0.25 ± 0.08 μm^2^/s; *D*_anti-His6 Ab_ = 0.18 ± 0.06 and 0.19 ± 0.06 μm^2^/s for imFCS and FRAP, respectively), concurring with the values obtained for the measurements upon addition of the nanotool. Analysis of clustered regions (1 μm) within the selected ROIs led to a further decrease in the diffusion coefficient (*D*_*spots*_= 0.15 ± 0.05 μm^2^/s and 0.13 ± 0.03 μm^2^/s for imFCS and FRAP, respectively). For imFCS measurements, two-sample t-tests (α = 0.05) were applied to compare the diffusion coefficients for the different conditions (***p≤ 0.001). The mean ± SD is shown. 36 and 26 cells for the conditions before and after addition of anti-His_6_ antibody were analyzed. For FRAP, the mean ± SD is shown. Here, 9 cells before, 5 cells after addition of anti-His_6_ antibody, 20x 1 μm ROIs were examined. **p≤ 0.01 for Tukey test. Scale bar: 1 μm.

**Figure 4–figure supplement 1.**
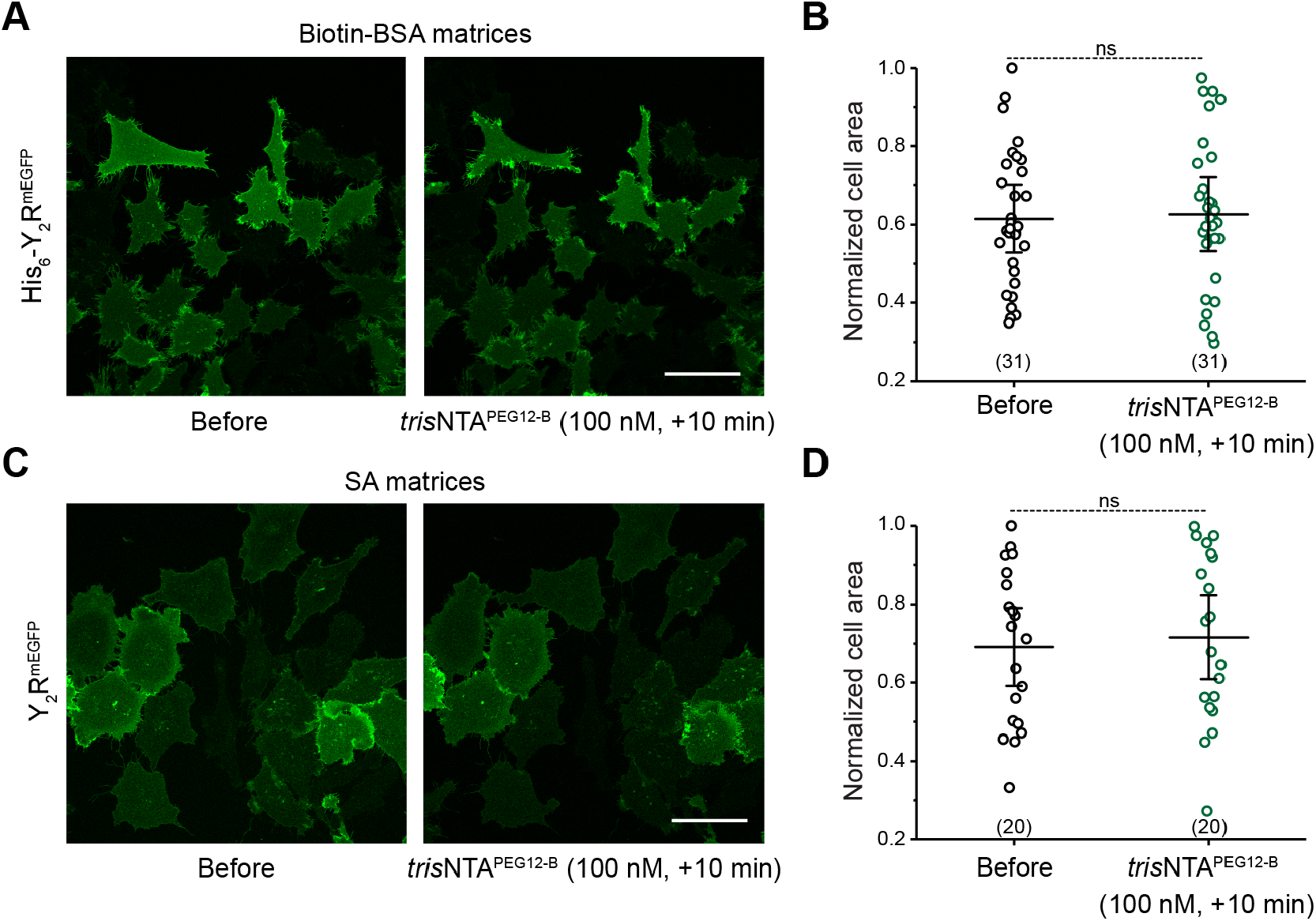
Changes in cell motility are exclusively triggered upon receptor clustering. (A, B) Confocal images of cells expressing His_6_-tagged Y_2_R on matrices which do not contain SA but biotin-BSA only. Addition of the *tris*NTA^PEG12-B^ nanotool confirmed no effect on cell spreading and motility as shown in the quantification of the cell area (B). The mean ± SD (31 cells) is shown. **p≤ 0.01 for Tukey test. (C) Confocal images of cells expressing Y_2_ receptors lacking the His_6_-tag on SA-matrices do not present significant changes in cells spreading upon addition of the nanotool. (D) Quantification of the cell area before and after addition of *tris*NTA^PEG12-B^ (100 nM). Values for cell area were normalized with respect to the highest value. The mean ± SD (20 cells) is shown. **p≤ 0.01 for Tukey test. Scale bar: 50 μm.

**Figure 5–figure supplement 1.**
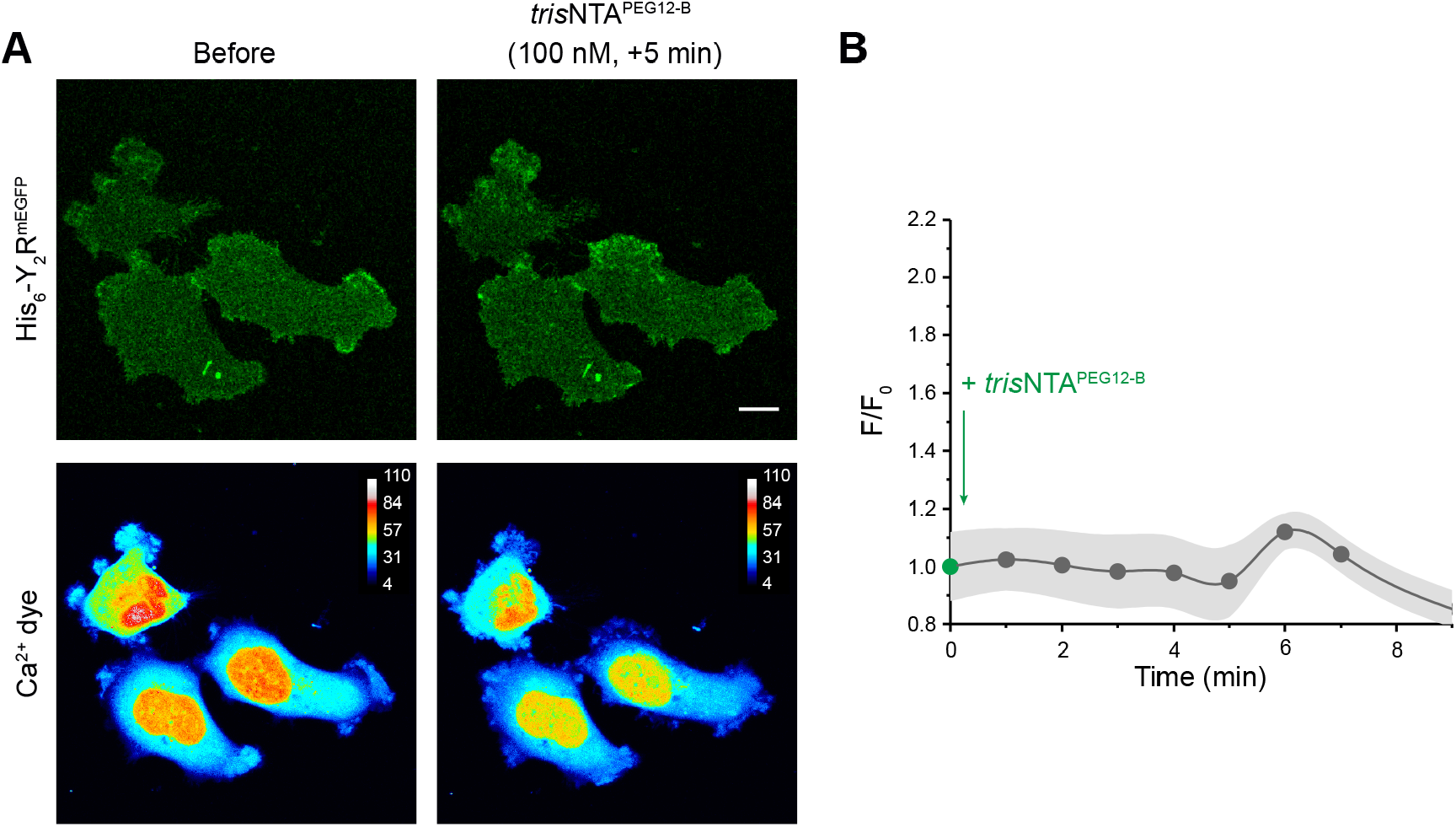
Calcium signaling is a specific response upon clustering. (A) Representative fluorescence images of the Y_2_R (upper panel) and color-coded images of the Ca^2+^ dye (lower panel). Y_2_R-expressing cells over pre-structured matrices in the absence of streptavidin were incubated with BioTracker 609 Red Ca^2+^ AM dye (3 μM) for 30 min. After rinsing, cells were immediately imaged by CLSM in LCIS at 37 °C. Addition of *tris*NTA^PEG12-B^ showed neither clustering nor change in cytosolic calcium. Scale bar: 10 μm. (B) Analysis of the mean gray value for Ca^2+^ signal before (F_0_) and upon (F) addition of *tris*NTA^PEG12-B^ *versus* time. Time-lapse images were recorded with 45 s interval before and after addition of *tris*NTA^PEG12-B^ (100 nM) (5 slices z-stack per time-point). ROIs covering the complete cell area were considered. The mean ± SD (10 cells) is shown.

**Figure 5–figure supplement 2.**
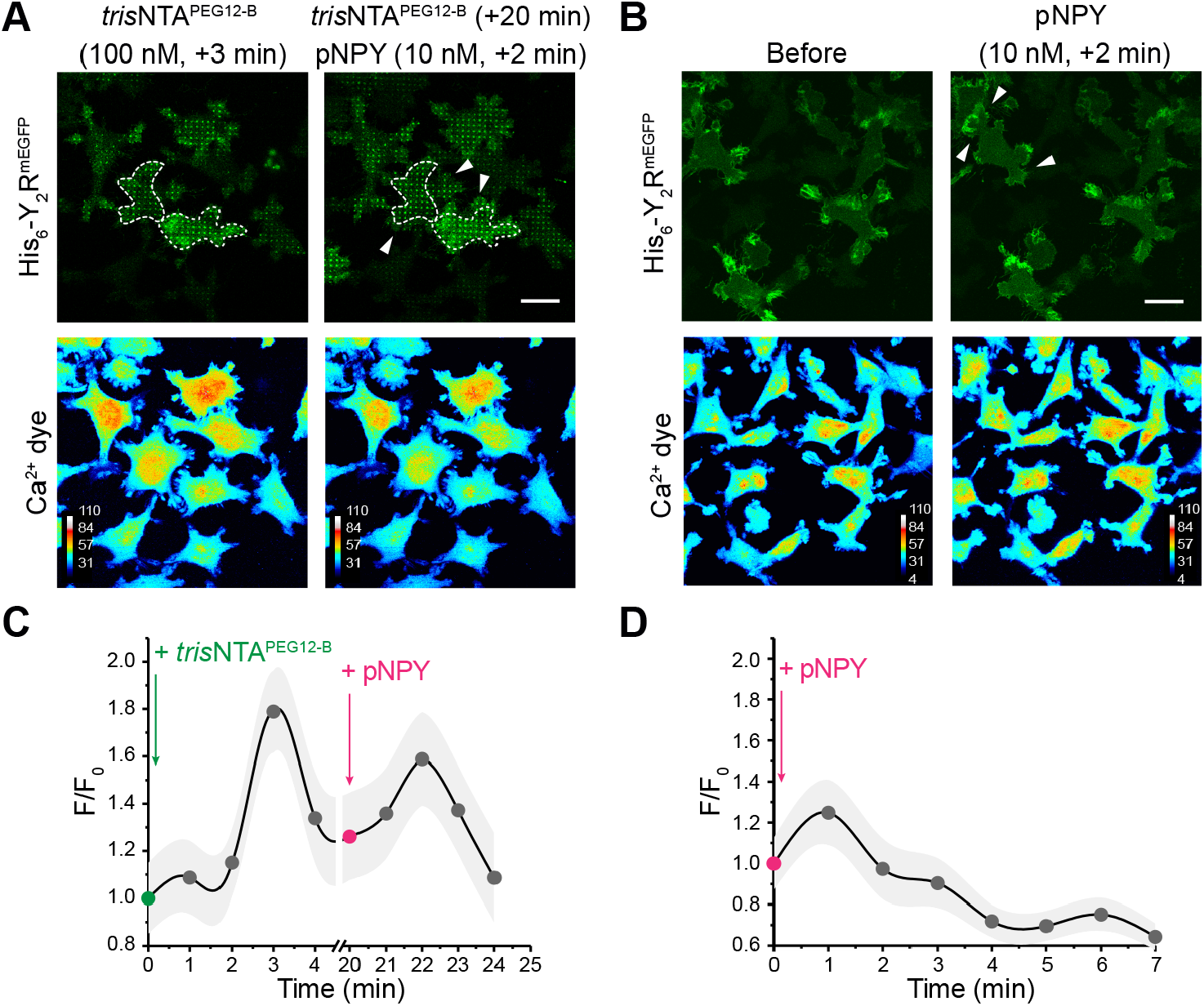
Receptor clustering potentiates calcium signaling. (A, B) Representative fluorescence images of the Y_2_R (upper panel) and color-coded images of the Ca^2+^ dye (lower panel). Y_2_R-expressing HeLa cells were allowed to adhere to pre-structured SA-matrices for 3 h. Before visualization, cells were incubated with BioTracker 609 Red Ca^2+^ AM dye (3 μM) for 30 min. After rinsing, cells were visualized by CLSM in LCIS at 37 °C and imaged before and after addition of only pNPY or before and after *tris*NTA^PEG12-B^ and subsequent addition of pNPY. Scale bar: 20 μm. (C, D) Analysis of the mean gray value for Ca^2+^ signal before (F_0_) and upon (F) addition of *tris*NTA^PEG12-B^/pNPY versus time. Time-lapse images were recorded with 45 s interval before and after addition of *tris*NTA^PEG12-B^/pNPY (5 slices z-stack per time-point). ROIs covering the complete cell area were considered. The mean ± SD (10 cells for each condition) is shown.

**Figure 6–figure supplement 1.**
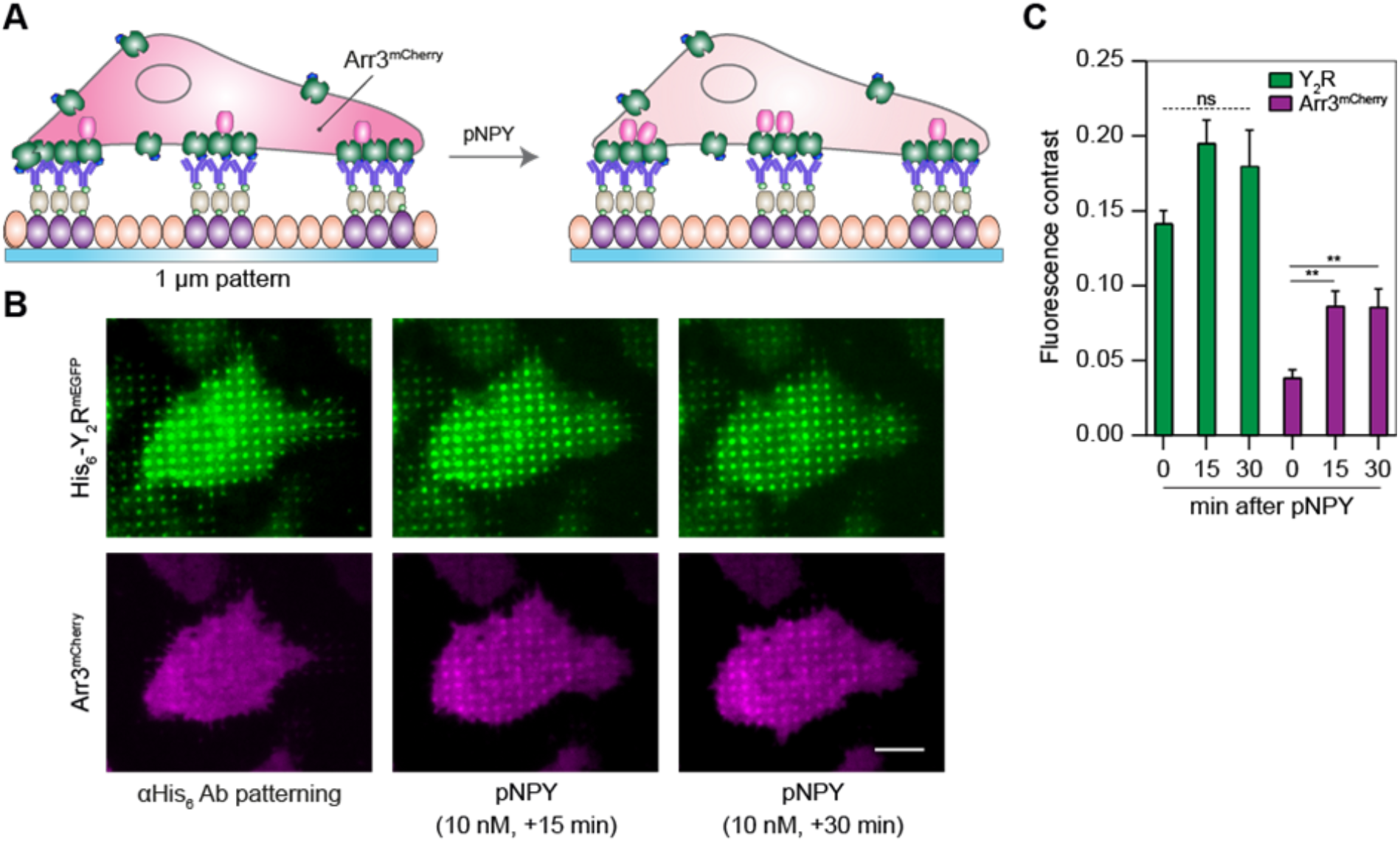
Arrestin-3 recruitment on antibody-confined regions. (A) Schematic representation of the experimental set-up. Cells co-expressing Y_2_R and Arr3 were allowed to adhere to anti-His_6_ antibody pre-structured matrices for 3 h and visualized by total internal reflection fluorescence (TIRF) microscopy in LCIS at 37 °C. (B) Representative TIRF images of cells before and upon addition of pNPY (10 nM) in LCIS for 30 min at 37 °C. Scale bar: 10 μm. (C) Quantification of the fluorescence contrast in the Y_2_R-patterned regions showed no significant change in receptor intensity yet a recruitment of Arr3 upon addition of pNPY (2-fold). Data are expressed as the means ± SEM (30 cells for each condition were analyzed). Tukey’s multiple comparison test was applied (**p≤ 0.01).

## Notes

### Competing Interest Statement

The authors have declared no competing interest.

